# Histone H3 Lysine 4 methyltransferases MLL3 and MLL4 Modulate Long-range Chromatin Interactions at Enhancers

**DOI:** 10.1101/110239

**Authors:** Jian Yan, Shi-An A Chen, Andrea Local, Tristin Liu, Yunjiang Qiu, Ah Young Lee, Inkyung Jung, Sebastian Preißl, Chloe M Rivera, Chaochen Wang, Haruhiko Ishii, Rongxin Fang, Zhen Ye, Kai Ge, Ming Hu, Bing Ren

**Affiliations:** Ludwig Institute for Cancer Research, 9500 Gilman Dr., La Jolla, CA 92093, USA; Department of Medical Biochemistry and Biophysics, Division of Functional Genomics and Systems Biology, Karolinska Institutet, 171 65 Stockholm, Sweden; Laboratory of Endocrinology and Receptor Biology, National Institute of Diabetes and Digestive and Kidney Diseases, NIH, Bethesda, MD 20892, USA; Department of Population Health, Division of Biostatistics, New York University, School of Medicine, 650 First Avenue, New York, NY 10016, USA; University of California San Diego, School of Medicine, Department of Cellular and Molecular Medicine, Institute of Genomic Medicine, 9500 Gilman Dr., La Jolla, CA 92093, USA

**Author notes:** These authors contributed equally (Jian Yan, Shi-An A Chen). Correspondence: Bing Ren.

## Abstract

Regulation of gene expression in mammalian cells depends on long-range chromatin interactions between enhancers and promoters. Currently, the exact mechanisms that connect distal enhancers to their specific target promoters remain to be fully elucidated. Here we show that the histone H3 Lysine 4 monomethylation (H3K4me1) writer proteins MLL3 and MLL4 (MLL3/4) play an active role in this process. We demonstrate that in differentiating mouse embryonic stem cells, MLL3/4-dependent deposition of H3K4me1 at enhancers correlates with increased levels of chromatin interactions, whereas loss of MLL3/4 leads to greatly reduced frequencies of chromatin interactions and failure of lineage-specific gene expression programs. We further show that H3K4me1 facilitates recruitment of the Cohesin complex to chromatin *in vitro* and *in vivo*, providing a potential mechanism for MLL3/4 to promote chromatin looping. Taken together, our results support an active role for MLL3/4 in modulating chromatin organization at enhancers in mammalian cells.

## INTRODUCTION

Enhancers play a critical role in regulating spatiotemporal gene expression programs in animals (Lee and Young, 2013; Levine, 2010; Levine et al., 2014; Makova and Hardison, 2015; Ong and Corces, 2011). These *cis*-regulatory elements recruit sequence-specific transcription factors (TFs) (Lee et al., 1993; Levine, 2010) and chromatin remodeling complexes to regulate target gene transcription from a distance (Dekker et al., 2013; Gorkin et al., 2014; Lee and Young, 2013; Levine et al., 2014; Nora et al., 2012; Ong and Corces, 2014). Active enhancers exhibit characteristic histone modifications such as H3K4me1 and H3K27ac, DNase I hypersensitivity, occupancy by H3.3 and H2A.Z histone variants, and production of short-lived RNA known as eRNAs (Aday et al., 2011; Buecker and Wysocka, 2012; Hardison and Taylor, 2012; Heintzman et al., 2009; Heintzman et al., 2007; Rajagopal et al., 2013; Roy et al., 2010). Based on these biochemical features, millions of candidate enhancers have been annotated in the human genome (Consortium, 2012; Kundaje et al., 2015).

Lineage-specific enhancers undergo step-wise activation during development, beginning with the binding of sequence-specific transcription factors, which recruit various chromatin remodeling complexes to promote chromatin modification and nucleosome dynamics (Ren and Yue, 2015). The chromatin modification H3K4me1 is a hallmark of the initial stage of enhancer activation, while the appearance of active chromatin marks such as H3K27ac defines an active state (Buecker and Wysocka, 2012; Ernst et al., 2011; Heintzman et al., 2009; Rada-Iglesias and Wysocka, 2011; Wang et al., 2015). *Mll3* and *Mll4* encode the histone H3 lysine 4 (H3K4) monomethytransferases with partially redundant functions (Hu et al., 2013; Lee et al., 2013; Wang et al., 2016a). Recruitment of MLL3 and MLL4 by transcription factors is necessary for enhancer-mediated activation of target genes during cellular differentiation (Herz et al., 2012; Hu et al., 2013; Lee et al., 2013). It has been reported that MLL3/4 regulate enhancer activation through the recruitment of the co-activator protein p300, a histone acetyltransferase that mediates H3K27 acetylation and transcriptional activation of target genes (Wang et al., 2016a).

Activation of enhancers is not only characterized by various histone modifications, but also accompanied by formation of long-range chromatin interactions between the distal *cis*-regulatory elements and the target gene promoters (Deng et al., 2012; Gorkin et al., 2014). Enhancer/promoter interactions are constrained in the topologically associating domains (TADs) (Symmons et al., 2014; Symmons et al., 2016), which are megabase-long chromosome regions characterized by highly frequent intra-domain chromatin interactions and infrequent inter-domain interactions (Dixon et al., 2012; Nora et al., 2012). TADs are highly conserved among different cell types and across species (Dixon et al., 2012; Schmitt et al., 2016). Disruption of the TAD boundaries can result in altered gene expression and developmental disorders (Andrey et al., 2013; Dixon et al., 2016; Schwarzer and Spitz, 2014). A number of molecules have been shown to regulate chromatin organization in mammalian cells. The Mediator and Cohesin Complexes have been reported to play an important role in enhancer/promoter interactions (Kagey et al., 2010). The insulator binding protein CTCF has also been implicated in the chromatin organization in mammalian cells (Gomez-Diaz and Corces, 2014; Narendra et al., 2015; Phillips-Cremins et al., 2013; Rao et al., 2014; Sanborn et al., 2015; Tang et al., 2015; Zuin et al., 2014). At the beta-globin locus, the transcription co-factor Ldb1 mediates long-range chromatin interactions between the locus-control-region (LCR) and target genes (Deng et al., 2012). Additionally, eRNAs have been proposed to promote enhancer/promoter interactions (Chapuy et al., 2013; Li et al., 2013). The Polycomb repressive complex PRC1 has been reported to maintain promoter-promoter interactions at silent developmental genes (Eskeland et al., 2010; Schoenfelder et al., 2015).

Recent chromatin contact maps generated in fly and mammalian cells have revealed a close connection between the chromatin modification state and the higher order chromatin architecture (Boettiger et al., 2016; Dixon et al., 2015; Jin et al., 2013; Sanyal et al., 2012; Sexton et al., 2012; Wang et al., 2016b). In particular, changes in local chromatin interactions have been shown to correlate with H3K4me1 dynamics during differentiation of human embryonic stem (ES) cells (Dixon et al., 2015). Based on these observations, we hypothesized that H3K4me1 monomethyltransferases MLL3 and MLL4 may modulate chromatin interactions at enhancers. To test this hypothesis, we examined chromatin interactions in mouse embryonic stem cells (mESCs) deficient for both MLL3 and MLL4 (DKO, hereafter). We observed that MLL3/4 are indeed required for maintaining proper chromatin interactions between enhancer and promoters in mESC. Through genome-wide analysis of chromatin contacts, we demonstrated that the local chromatin contacts at distal regulatory elements are dependent on MLL3/4 in mouse ES cells. We obtained evidence that MLL3/4 modulate local chromatin organization through recruitment of the Cohesin complex. In the absence of MLL3/4, Cohesin occupancy at enhancers was greatly reduced during mES cell differentiation. Additionally, targeted deposition of H3K4me1 was sufficient for recruitment of Cohesin complex to DNA in cells. Altogether, these results provided strong evidence that MLL3/4 activate enhancers by promoting chromatin interactions, in addition to recruitment of p300.

## RESULTS

### MLL3/4 are required for enhancer/promoter chromatin interactions at *Sox2*

To determine the role of MLL3/4 on chromatin organization at enhancers, we first focused on a previously identified super-enhancer located 130 kb downstream of the *Sox2* gene (Figure 1A and **Supplemental** Figure S1A) (Li et al., 2014; Zhou et al., 2014). Deletion of this super-enhancer caused a drastic, allele-specific reduction of Sox2 expression in *cis* (Li et al., 2014; Zhou et al., 2014). Additionally, this super-enhancer was shown to interact with the *Sox2* gene in mESCs (Li et al., 2014; Phillips-Cremins et al., 2013; Zhou et al., 2014). Both H3K4me1 and H3K27ac were present at the super-enhancer in wild-type cells (WT, hereafter), but were completely or partially lost in MLL3/4-knockout cells (DKO, hereafter) (Figure 1A), confirming an essential role for MLL3/4 in deposition of active chromatin marks at the *Sox2* distal super enhancer (*Sox2* SE, hereafter).

**Figure 1.**
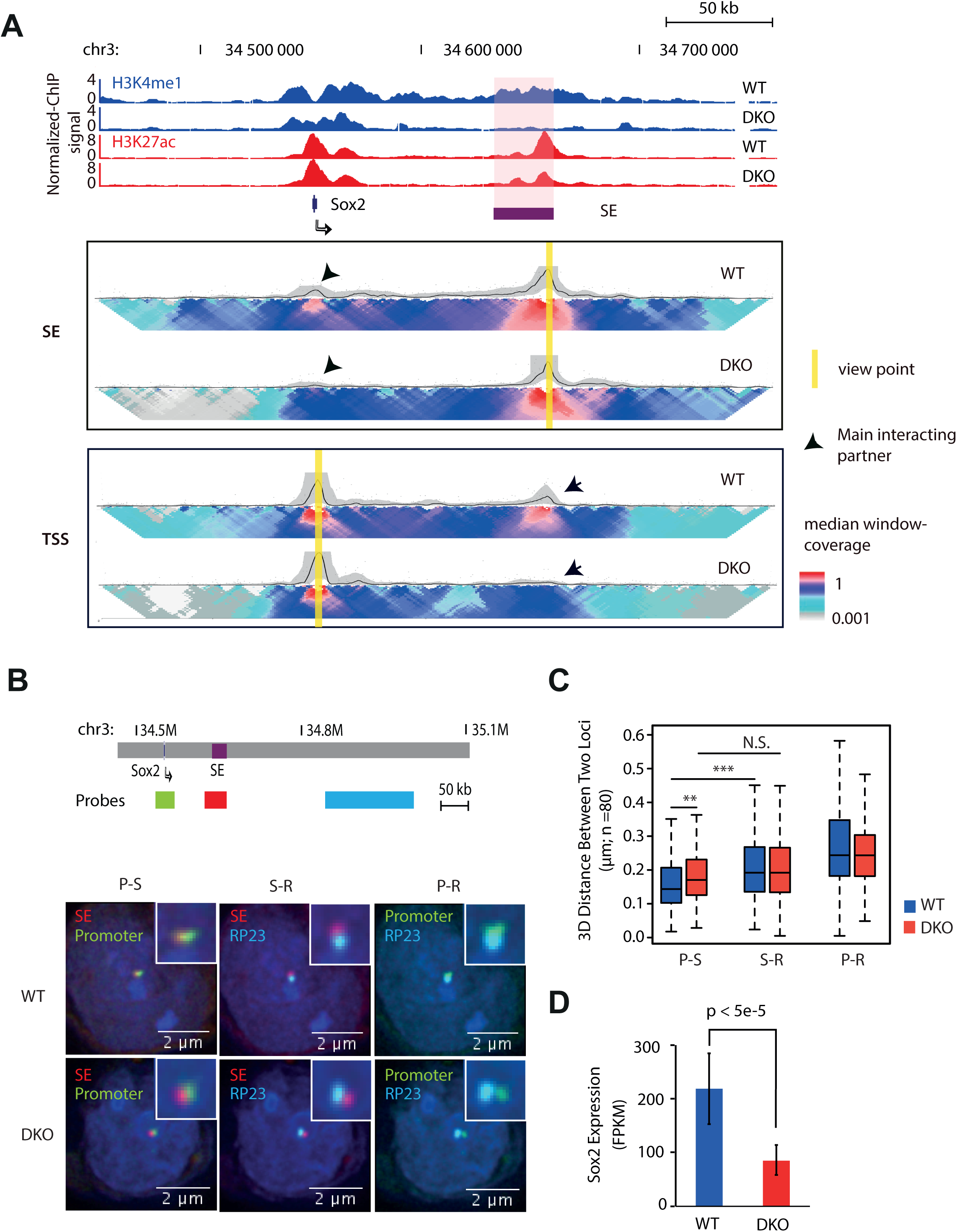
MLL3/4 Are Required for Chromatin Interactions at the *Sox2* enhancer. **(A)** ChIP-seq and 4C-Seq analysis of *Sox2* locus in wild type and MLL3/4 double knockout mESCs. Top, genome browser snapshot of ChIP-seq data showing loss of H3K4me1 and partial reduction of H3K27Ac at *Sox2* Super-enhancer (SE) in MLL3/4 double knockout mESCs. SE, super-enhancer. WT, wild type mouse ES cell line E14. DKO, MLL3/4 double knockout mouse ES cell line. y-axis shows input normalized ChIP-seq reads per kilobase pair per million total reads (RPKM). *Sox2* SE is indicated in red shade. *Sox2* gene locus is indicated by arrow. Bottom, 2D-heat map of 4C-seq analysis showing significant reduction in contact frequency between *Sox2* TSS and *Sox2*-SE in DKO cells relative to WT cells. The same genomic position is aligned with genome browser snapshot for ChIP-seq analysis. Panel SE, 4C-seq with viewpoint at *Sox2* super-enhancer locus, highlighted by yellow bar. Panel TSS, 4C-seq viewpoint at *Sox2* TSS locus, highlighted by yellow bar. Black arrows emphasize the main interacting partner with the viewpoint. Heatmap shows the median genomic coverage using different sizes of sliding windows between 2 kb (top row) and 50 kb (bottom row). The gray shade above the heatmap shows the genomic coverage between 20th and 80th quantile values using a slide window of 5 kb. The black trend line indicates the median genomic coverage using a slide window of 5 kb. **(B)** 3D FISH microscopy images showing that physical distance between *Sox2* SE and promoter becomes larger in DKO mESC, relative to WT. Top, schematic showing relative genomic positions for 3D FISH probes. Bottom, 3D FISH images showing overlay signals for probe set combinations in representative WT and DKO mESC. Nucleus was stained with DAPI. Insets show the zoom in of probe-detected foci for clearance. Red color, probes hybrid to SE locus, Green color, probes detecting promoter locus; Cyan color, probes detecting a region (RP23) that is located 100 kb downstream from SE. Note that Promoter is approx. 100 kb upstream of SE. **(C)** Summary of FISH data from approximately 80 individual cells for both WT and DKO cell types. y-axis shows the distance between the centers of two foci represented by two colors. Scale bar is shown at the bottom right corner of each image. Note that distance between promoter and SE was significantly larger in DKO than WT. S-R, distance between RP23 and SE; P-R, distance between RP23 and Promoter; P-S, distance between Promoter and SE. Stars indicate statistical significance tested with Mann-Whitney-U test (^**<^0.01;^***^ p^<^0.005). **(D)** Expression of Sox2 decreased upon *Mll3/4* DKO. FPKM is shown in y-axis. Error bars are derived from three biological replicates. See also Figure S1, Figure S2

To test the hypothesis that loss of MLL3/4 would impair chromatin interactions at enhancers, we employed 4C-seq (van de Werken et al., 2012) to examine the chromatin interactions centered at either the *Sox2* promoter or the distal enhancers, in WT cells and DKO cells (Figure 1A). Strikingly, the *Sox2* promoter-enhancer interaction was dramatically reduced in the DKO cells, suggesting that MLL3/4 are required for the formation of chromatin interactions between the *Sox2* promoter and *Sox2* SE (Figure 1A and **Supplemental** Figure S1B). As a control, we examined a homozygous *Sox2* SE deletion mouse ES cell line (DEL, hereafter)(Li et al., 2014). Loss of interaction between *Sox2* promoter and *Sox2* SE flanking regions was detected in DEL cells, confirming that the super-enhancer element required for promoter-distal enhancer interactions at the *Sox2* locus (**Supplemental** Figure S1C). We further employed Three-Dimensional Fluorescent *in situ* Hybridization technology (3D-FISH) as an orthogonal approach to validate the loss of interaction *in vivo* at single cell resolution. Consistent with 4C-seq results, the spatial distance between *Sox2* SE and *Sox2* promoter significantly increased in both DKO and DEL cells (Figure 1B, **1C** and **Supplemental** Figure S1D, **S1E**).

In accord with previous observations linking *Sox2* promoter-enhancer interactions to *Sox2* transcription (Li et al., 2014; Zhou et al., 2014), a partial decrease in SOX2 mRNA expression was observed (~50%) in DKO cells (Figure 1D). Previously, it was shown that removal of *Sox2* SE led to more than 90% reduction of SOX2 expression (Li et al., 2014). The relative mild effect of MLL3/4 knockout on SOX2 expression indicates that additional mechanisms may be involved in regulating SOX2 expression that compensate for the reduction of MLL3/4 at the enhancer and subsequent loss of long-range interactions.

To exclude the possibility that loss of chromatin interactions in DKO cells is due to a reduced level of SOX2 expression, we carried out RNA-seq analysis of the WT and DKO cells. No apparent change in expression of other pluripotency transcription factors (TFs), such as Pou5f1 and Nanog, was detected (**Supplemental** Figure S1G, Figure 4B). Therefore, the loss of chromatin interactions between the super-enhancer and target gene in the DKO cells is unlikely the result of secondary effects of reduced expression or other transcription factors.

Similar to the *Sox2* locus, we also observed disruption of enhancer-promoter interactions and target gene expression at the *Car2* gene locus in DKO cells (**Supplemental** Figure S1F), while an independent study identified a loss of promoter/enhancer interactions at the *Lefty* gene locus using the same MLL3/4 DKO cells (Wang et al., 2016a). Our results demonstrate that chromatin interactions between enhancers and target genes depend on H3K4me1-methyltransferases MLL3/4.

### MLL3/4 loss leads to reduced chromatin interactions at enhancers in the ES cells

To determine whether MLL3/4 knockout also led to a general loss of chromatin interactions at enhancers in the DKO cells, we first carried out ChIP-seq analysis in WT and DKO cells to examine the effects of MLL3/4 loss on genomic distribution of H3K4me1. Focusing on a set of enhancers previously determined in mouse ES cells (Hnisz et al., 2013), we found that loss of MLL3/4 led to a significant decrease of H3K4me1 signals at enhancers genome-wide (**Supplemental** Figure S2A). We identified a total of 78,645 narrow H3K4me1 peaks in WT mESC cells, and observed that H3K4me1 signals at 34,527 of them decreased by more than 50% in DKO cells. We referred to these as MLL3/4-dependent H3K4me1 regions (**Supplemental** Figure S2B and **Table S1A**). Little or no change was detected in the other 44,118 genomic regions, which we designated as MLL3/4-independent H3K4me1 regions (**Supplemental** Figure S2B and **Table S1B**). In addition, 9,228 genomic regions acquired H3K4me1 peaks upon MLL3/4 knockout (Other; **Supplemental** Figure S2B and **Table S1C**). Consistent with a previous report (Hu et al., 2013), we found that more than 85% of MLL3/4-dependent H3K4me1 regions were promoter-distal (>2kb), whereas MLL3/4-independent H3K4me1 regions were enriched at or near TSS. Most of the increased H3K4me1 peaks in DKO were located within TSS proximal regions (**Supplemental** Figure S2C). Motif analysis revealed strong enrichment of ES-specific transcription factor (TF) binding sites in MLL3/4-dependent H3K4me1 regions (**Supplemental** Figure S2D), consistent with a previously reported role for the sequence-specific TFs in recruiting MLL3/4 to establish H3K4me1 at distal enhancers (Lee et al., 2013). Gene ontology analysis indicates that genes near the MLL3/4-dependent H3K4me1 regions are involved in pluripotency and stem cell maintenance (**Supplemental** Figure S2E), suggesting a role for MLL3/4-dependent H3K4me1 regions in cell type specific gene expression.

We next employed *in situ* Hi-C (Rao et al., 2014) to investigate alterations in chromatin architecture in both DKO cells to determine whether loss of MLL3/4 would result in loss of chromatin interactions at enhancers genome-wide. We carried out two biological replicates both WT and DKO cells, and obtained approximately 10^9^ reads from each cell line. As shown in Figure 2A, the TAD structures were well preserved upon MLL3/4 knockout (Figure 2A and **Supplemental** Figure S3A). The difference of TAD boundaries between WT and DKO is at a similar level of that between biological replicates (**Supplemental** Figure S3C and S3D). The strongest loss of contact frequency was observed between bins located shorter than 100 kb apart from each other (**Supplemental** Figure S3B). The short distance suggests that the effects of MLL3/4 loss on chromatin interactions are constrained within TADs. Since chromatin interactions that were altered between genomic regions were generally less than 200 kb apart, we focused on such local chromatin interactions in subsequent analyses. We observed that these short-range chromatin contacts were not evenly distributed in the genome. Some regions showed significantly higher frequencies of interactions than the others, which we referred to as Frequently Interacting REgions, or FIREs in short (Schmitt et al., 2016). To quantify local chromatin contacts, we calculated the accumulated contact frequency of a given bin to all its surrounded bins with a distance between 15 kb to 200 kb. As we focus on this specific range of interactions, we used the HiCNormCis for normalization (Schmitt et al., 2016, **Extended Experimental Procedures**), which achieved greater reduction or even complete elimination of systematic errors introduced by the effective fragment length, GC content and mappability score, than other normalization tools (**Supplemental** Figure S3E, **S3F**, **S3G** and **S3H**) (Imakaev et al., 2012; Rao et al., 2014). We defined FIRE score as the z-score for each bin. We found that FIRE scores were highly reproducible between biological replicates (Pearson r_WT_=0.964, **Supplemental** Figure S4A; and r_DKO_=0.960, **Supplemental** Figure S4B), but were significantly different between the WT and DKO cells (**Supplemental** Figure S4C). This result suggests that FIRE score is a robust and sensitive proxy for local chromatin interactions including promoter-enhancer interactions. By using a threshold of p value <= 0.05, we determined 14,190 FIREs in WT cells and 13,542 FIREs in DKO cells, covering approximately 5% of total genome. Roughly 70% of FIREs are shared between WT and DKO cells. We categorize FIREs into WT-specific, DKO-specific and shared, with a cutoff of FIRE score = 1.5 (**Supplemental** Figure S4D, **Table S2**). Genes in the cell-type (WT or DKO) specific FIREs showed significantly higher expression level than the same set of genes expressed in the other cell type (**Supplemental** Figure S4E). WT-specific FIREs were significantly enriched in genes involved in regulation of cell differentiation and cell fate determination (**Supplemental** Figure S4F).

**Figure 2.**
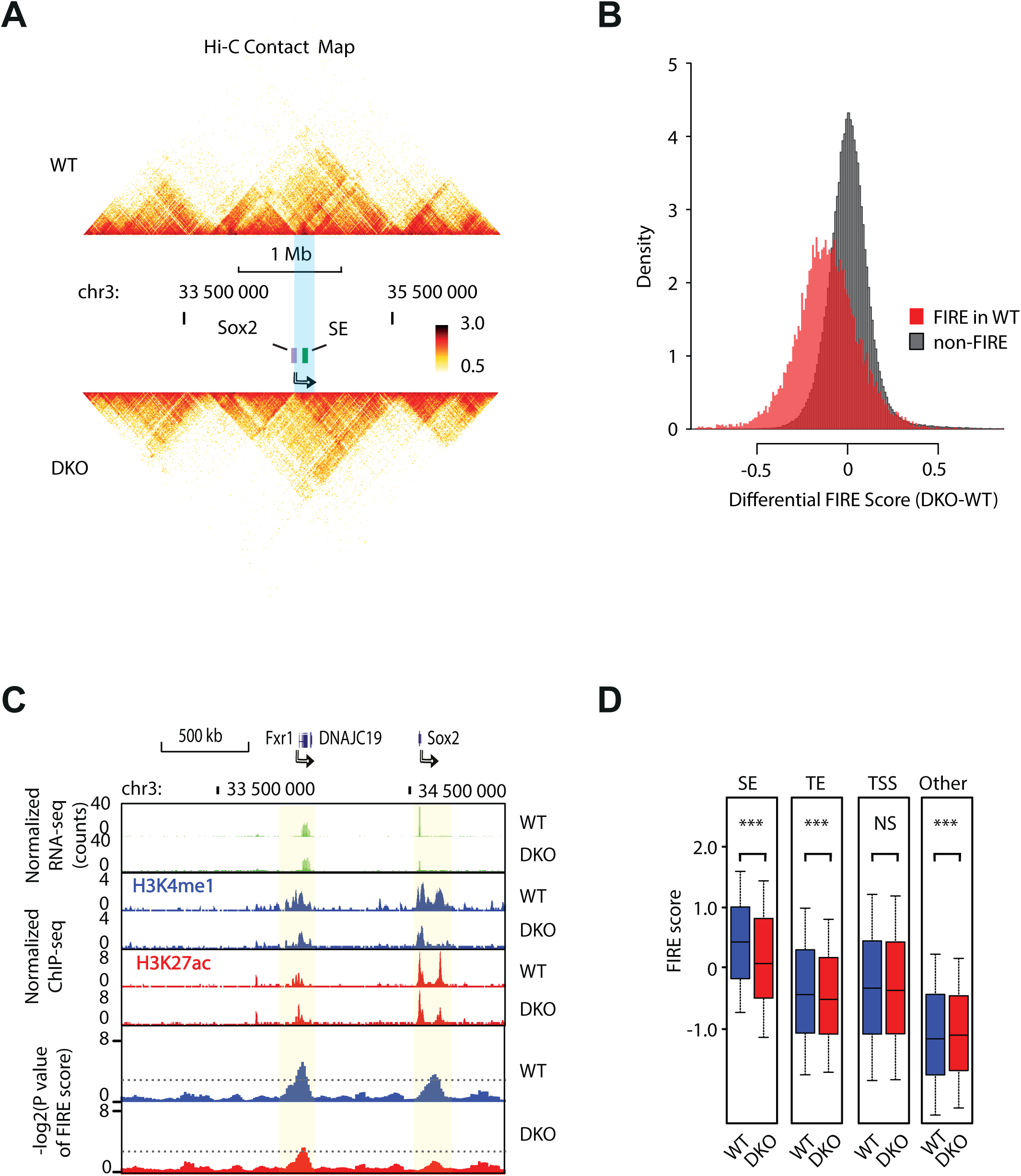
MLL3/4-dependent H3K4me1 Show Reduced Chromatin Interactions in DKOmESC. **(A)** Heatmap showing the chromosomal contacts near the *Sox2* locus, as determined in *in situ* HiC experiments. The upper panel shows the WT cells and lower panel shows the DKO cells. *Sox2* gene is indicated by the violet box and an arrow. Color key shows the normalized contact frequency. **(B)** Histogram showing the distribution of differential FIRE scores across 10-kb bins (DKO-WT). Red, bins classified as FIRE regions in WT cells. Grey, non-FIRE bins in WT cells. **(C)** Genome browser track shows correlation between the change of FIRE scores (bottom) associated and changes of H3K4me1 ChIP-seq RPKM (middle) upon MLL3/4 knockout. Partial loss of SOX2 expression is also shown (top). For comparison, FXR1 or DNAJC19 expression is stable, consistent with the stable FIRE structure. The light green shades indicate the FIRE regions. Dashed lines label the cutoff for FIRE. **(D)** Boxplots comparing FIRE scores in WT (blue) and DKO (red) for bins containing different classes of *cis*-regulatory elements (TE, SE and TSS). SE, super-enhancer. TE, typical enhancer. TSS, transcription starting site. Other, bins with increased FIRE scores in DKO relative to WT. The p value below each category was computed by two-tail paired Welch t-test. Asterisk indicates that the difference is statistically significant. Note that SE displays the highest FIRE score among all elements tested here and SE, TE and TSS generally have higher FIRE score than the average genome. See also **Table S1**, Figure S3, Figure S4

Upon knockout of MLL3/4, the chromatin contacts in WT FIREs were reduced compared to non-FIRE genomic regions (Figure 2B). To illustrate how H3K4me1 signals are associated with chromatin interactions at individual loci, we used a 2 Mb-region containing *Sox2* on chromosome 3 as an example. The FIRE score was significantly decreased near the *Sox2* SE locus in DKO cells, coincident with depletion of H3K4me1, whereas the nearby *Fxr1* locus was less affected (Figure 2C). We observe a significant decrease in average FIRE scores of typical (TE) and super enhancers (SE) genome-wide, but not at the transcriptional start sites (TSS) **(Figure 2D)**. Consistent with this observation, FIRE scores at WT-specific H3K4me1 peaks were higher in WT cells compared to DKO cells (**Supplemental** Figure S4G). Similarly, WT-specific and shared H3K27ac peaks between WT and DKO showed average increase in FIRE score (**Supplemental** Figure S4H). The FIRE score, a well-established proxy for chromatin interactions (Schmitt et al., 2016), was demonstrated here tightly associated with MLL3/4-dependent H3K4me1 peaks and H3K27ac peaks. These peaks that are mostly located at TSS distal enhancer regions (**Supplemental** Figure S2). Our results suggest that MLL3/4 are required for elevated levels of chromatin interactions at the enhancers compared with other genomic regions.

### The Cohesin Complex Acts Downstream of MLL3/4 to Promote Chromatin Interactions at Enhancers

To gain insight into the molecular mechanisms by which MLL3/4 promote chromatin interactions at enhancers, we next examined the inter-dependency of enhancer occupancy by MLL3/4 and Cohesin complex (Kagey et al., 2010). The Cohesin complex, consisting of Smc1, Smc3, Rad21 and SA1/2, has been proposed to mediate chromatin contacts between enhancers and promoters (Hadjur et al., 2009; Ing-Simmons et al., 2015; Kagey et al., 2010; Mizuguchi et al., 2014; Wendt et al., 2008; Yan et al., 2013; Zuin et al., 2014). We hypothesized that MLL3/4 may facilitate the recruitment of Cohesin complex to enhancers. To test this hypothesis, we first asked whether MLL3/4 are required for the recruitment of Cohesin at the *Sox2* SE. We performed ChIP-seq experiments in WT and DKO cells using antibodies against the Cohesin subunit Rad21 and Mediator subunit Med12. Consistent with previous reports (Kagey et al., 2010; Yan et al., 2013), both Cohesin and Mediator complexes were co-localized with H3K4me1 peaks at the *Sox2* Super-enhancer in WT mESC (Figure 3A). In addition, we observed elevated FIRE scores at the Cohesin and Mediator binding sites, consistent with previous report that Cohesin act together with the Mediator complex to mediate chromatin interactions (Kagey et al., 2010)(Figure 3F). Upon knockout of MLL3/4, occupancy of the *Sox2* Super-enhancer by both complexes was lost or greatly reduced in WT and DKO cells (Figure 3A), suggesting that occupancy of Cohesin and Mediator complexes at enhancers is MLL3/4-dependent. Genome-wide analysis further showed that binding of Cohesin complex to MLL3/4-dependent H3K4me1 peaks was drastically reduced, while the occupancy remained unchanged near the *Mll3/4*-independent H3K4me1 peaks (Figure 3B).

**Figure 3.**
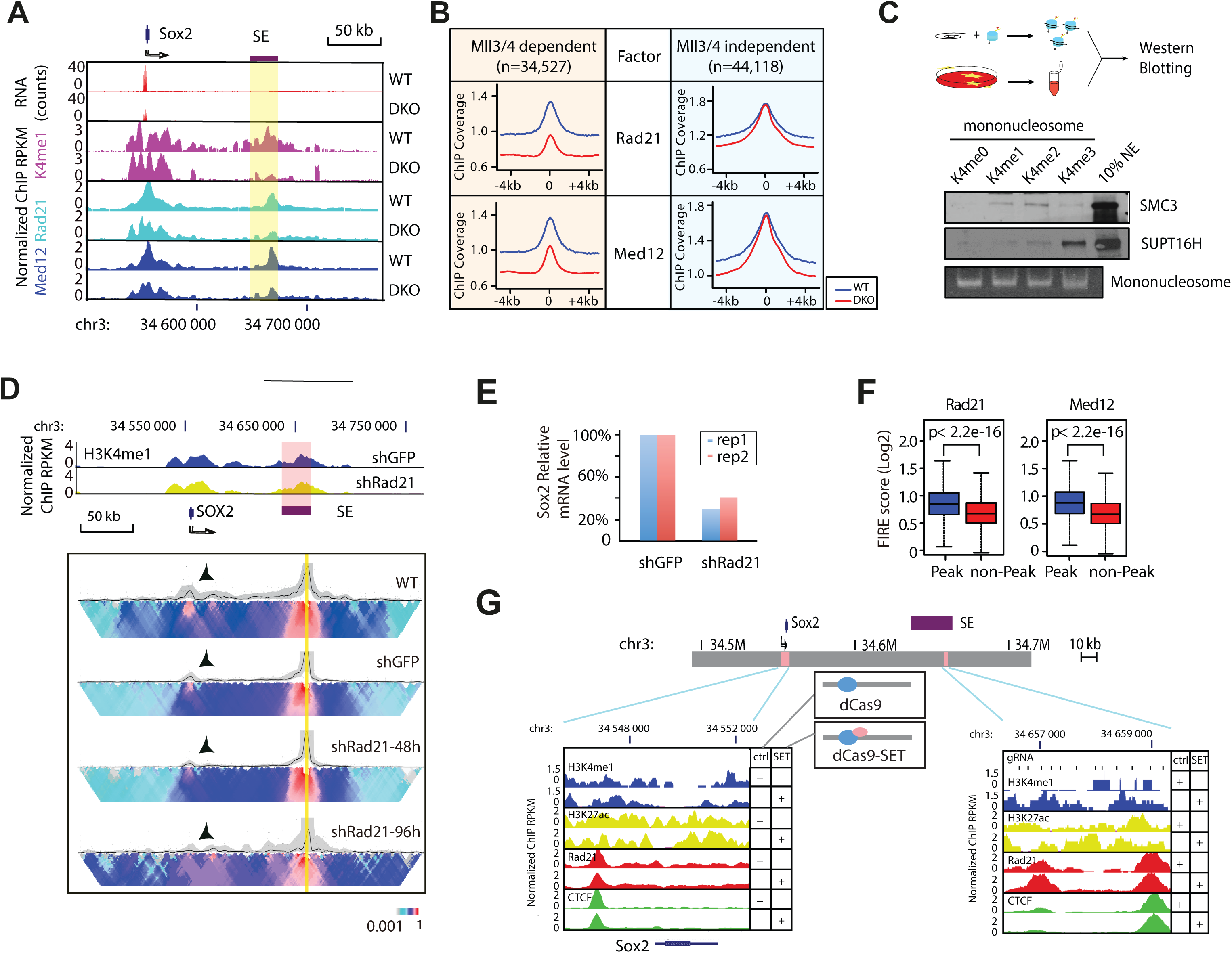
MLL3/4-Dependent H3K4me1 Facilitates Cohesin Binding to Chromatin. **(A)** Genome browser tracks showing loss of Cohesin (Rad21) and Mediator (Med12) binding at the *Sox2* gene and *Sox2* super-enhancer (SE, indicated in violet shade) in MLL3/4 DKO cells. Top, relative genomic positions of the *Sox2* gene and *Sox2* SE. Middle, RNA-seq tracks show Sox2 expression for reference, quantified in **Figure 1D**. Bottom, Normalized ChIP-seq signals of H3K4me1, Rad21 and Med12 at the *Sox2* locus. **(B)** ChIP-seq analysis showing binding of Cohesin and Mediator complexes to MLL3/4-dependent H3K4me1 peaks. ChIP-seq signals are centered around H3K4me1 peaks and extended 4kb upstream and downstream along the genome. X-axis indicates relative coordinates to peaks center. Y-axis indicates z-score of input-normalized ChIP-seq RPKM value. Blue, aggregated ChIP-seq signal in WT cells. Red, aggregated ChIP signal in DKO cells. Note that Cohesin (Rad21) and Mediator (Med12) are affected at MLL3/4-dependent H3K4me1 regions in MLL3/4 DKO cells, coinciding with the loss of H3K4me1. **(C)** The *in vitro* pull-down assay showing that Cohesin (Smc3) preferentially associate to H3K4me1 and H3K4me2 mononucleosomes. Top, schematic showing assay workflow. Modified H3 histones are assembled into biotinylated DNA-bound nucleosomes *in vitro*. Nucleosomes were then incubated with HeLa nuclear lysate and the Streptavidin pull-down fractions were assayed for binding factors with Western blotting. Bottom, Western blots showing binding of SMC3 and SUPTH binding to H3K4 unmodified (K4me0), mono-(K4me1), di-(K4me2) and tri-(K4me3) methylated nucleosomes. The FACT complex subunit SUPT16H is used as a control to show that FACT binds preferentially to H3K4me3 mono-nucleosomes. Agarose gel staining for 601λ DNA was used as loading control for mononucleosomes. **(D)** Genome browser tracks showing knockdown of Cohesin complex (shRad21) does not cause overt change of H3K4me1 signals and 4C interactions at the *Sox2* locus. Top, Normalized H3K4me1 ChIP-seq tracks for control and Rad21 knockdown cells. shGFP, control using shRNA against GFP sequences. SE, super-enhancer, indicated also by violet shade. Bottom, 4C-seq analysis showing that chromatin interactions between *Sox2* SE and promoter are reduced upon Rad21 depletion by shRNA. shRad21-48h, shRNA knockdown targeting RAD21 mRNA 48 hours after lentivirus infection. shRad21-96h, shRNA knockdown targeting RAD21 mRNA 96 hours after lentivirus infection. Black arrows emphasize the main interacting partner with the viewpoint. Heatmap shows the median genomic coverage of the indicated position using different sizes of sliding windows between 2 kb (top row) and 50 kb (bottom row). The gray shade above the heatmap shows the genomic coverage between 20th and 80th quantile values using a slide window of 5 kb. The black trend line indicates the median genomic coverage using a slide window of 5 kb. Note that the interaction between Sox2 SE and gene body is dramatically lost at 96 hours post knock-down via lentivirus. **(E)** Bar chart showing reduction of Sox2 expression upon knockdown of RAD21 using shRNA in two biological replicate experiments. *Sox2* mRNA levels were quantified by RT-qPCR and normalized to β-Actin mRNA levels. Y-axis, Sox2 mRNA levels relative to shGFP. Blue, first replicate. Red, second replicate. **(F)** Box plots showing FIRE scores for 10-kb bins that include and exclude Rad21 or Med12 peaks. Note that both Rad21 and Med12 including bins show higher FIRE score over non-including bins in average. p values are computed with paired t-tests. **(G)** Genome browser tracks showing increased H3K4me1 and Cohesin (RAD21) binding in the *Sox2* SE locus in DKO cells expressing dCas9-MLL3SET. DKO cells were transfected with vectors co-expressing tiling CRIPSR guides targeting *Sox2* SE and dCas9 proteins with or without MLL3SET domain fusion. Top, schematic of the loci assayed. Pink boxes indicate genomic regions of interest. SE(purple), *Sox2* super enhancer. Bottom panels, genome browser tracks showing normalized ChIP-seq coverage in RPKM for H3K4me1, H3K27ac, RAD21 and CTCF at control region (*Sox2* promoter) and the guide RNA targeted region of *Sox2* SE. Y-axis, relative RPKM coverage was calculated by normalizing to the constant, CTCF-overlapping major peak right to the dashed box. Ctrl: dCas9 without MLL3SET domain. SET: dCas9-MLL3SET fusion protein. See also Figure S5

To determine whether Cohesin complex acts downstream of MLL3/4 to regulate chromatin interactions, we depleted the expression of Cohesin complex component RAD21 using shRNA-mediated knockdown (**Supplemental** Figure S5C). On the other hand, no obvious change of H3K4me1 at either *Sox2* SE or around the gene body was detected (Figure 3D, **upper panel**), despite loss of chromatin interactions between *Sox2* SE and the promoter (Figure 3D, **lower panel**). *Sox2* expression also decreased upon *Rad21* knockdown, confirming the functional role of enhancer-promoter interactions in gene activation (Figure 3E). Therefore, MLL3/4 likely act upstream of the Cohesin complex to mediate chromatin interactions.

### H3K4me1 Facilitates Recruitment of Cohesin Complex

One potential mechanism for MLL3/4 to facilitate recruitment of Cohesin complex at enhancers is via H3K4me1. Consistent with this hypothesis, the Cohesin complex is generally co-localized with H3K4me1 peaks *in vivo* (Figure 3B and **Supplemental** Figure S5B). To further test this hypothesis, we performed *in vitro* pull down assays using nuclear extracts from HeLa cells and reconstituted mononucleosomes bearing either unmodified H3 histones or H3 with chemically-modified epitopes mimicking lysine 4, including mono-, di- and tri-methylation. We found that the Cohesin complex bound more strongly to the mononucleosomes with H3K4me1 and H3K4me2 modifications than unmodified nucleosomes. As a control, the FACT complex showed stronger preference to H3K4me3 than H3K4me1 (Orphanides et al., 1999; Takahata et al., 2009)(Figure 3C and **Supplemental** Figure S5A).

Our ChIP-seq and *in vitro* pull-down assays strongly suggest that H3K4me1 may either directly or indirectly facilitate the binding of the Cohesin complex to enhancers to mediate chromatin interactions. To further determine whether H3K4me1 is the sufficient for Cohesin recruitment, we repurposed the catalytically dead Cas9 protein (dCas9) to induce ectopic H3K4me1 at targeted locus by fusing it to the MLL3 SET domain (MLL3SET, hereafter), which catalyzes monomethylation of H3K4 (Hu et al., 2013). We show that transfection of dCas9-MLL3SET and a set of guide RNAs targeting the Sox2 SE to DKO cells significantly increases local H3K4me1 levels at the targeted region (Figure 3G, **right panel**) as well as Cohesin occupancy at Sox2 SE, indicating that H3K4me1 ca indeed facilitate Cohesin recruitment in vivo. As a control, H3K4me1 and Cohesin occupancy at the non-targeted *Sox2* promoter was not significantly altered (Figure 3G, **left panel**).

### Dynamic Chromatin Organization at Lineage-specific Enhancers during Mouse Stem Cell Differentiation

If MLL3/4 plays an important role in chromatin interactions at enhancers, its presence at enhancers during mESC differentiation would be required for new chromatin interactions at enhancers to occur. To test this prediction, we treated WT and DKO cells with retinoic acid (RA) to induce the differentiation of these cells towards the neural progenitor cell (NPC) lineage. We collected the cells every 12 hours over a 60-hour period (Figure 4A). Single cell RNA-seq analyses showed that both WT and DKO cells lost Nanog expression after RA treatment. However, while the WT cells successfully differentiate into the NPC lineage, as evidenced by the expression of NPC-specific markers, such as Vimentin, while DKO cells failed to completely differentiate into the NPC lineage (Figure 4B). Bulk RNA-seq analysis showed broad defects in the induction of genes involved in neuronal function (Figure 4C, D). In particular, group III genes were induced in WT but not in DKO cells (Figure 4C). Lack of induction of these genes in DKO cells indicated a failure of differentiation towards NPC in the absence of MLL3/4, supported by Gene Ontology analysis of group III genes (Figure 4D). This result strongly supports the model that MLL3/4 are critical for stem cell differentiation and renewal (Eissenberg and Shilatifard, 2010; Gu and Lee, 2013).

**Figure 4.**
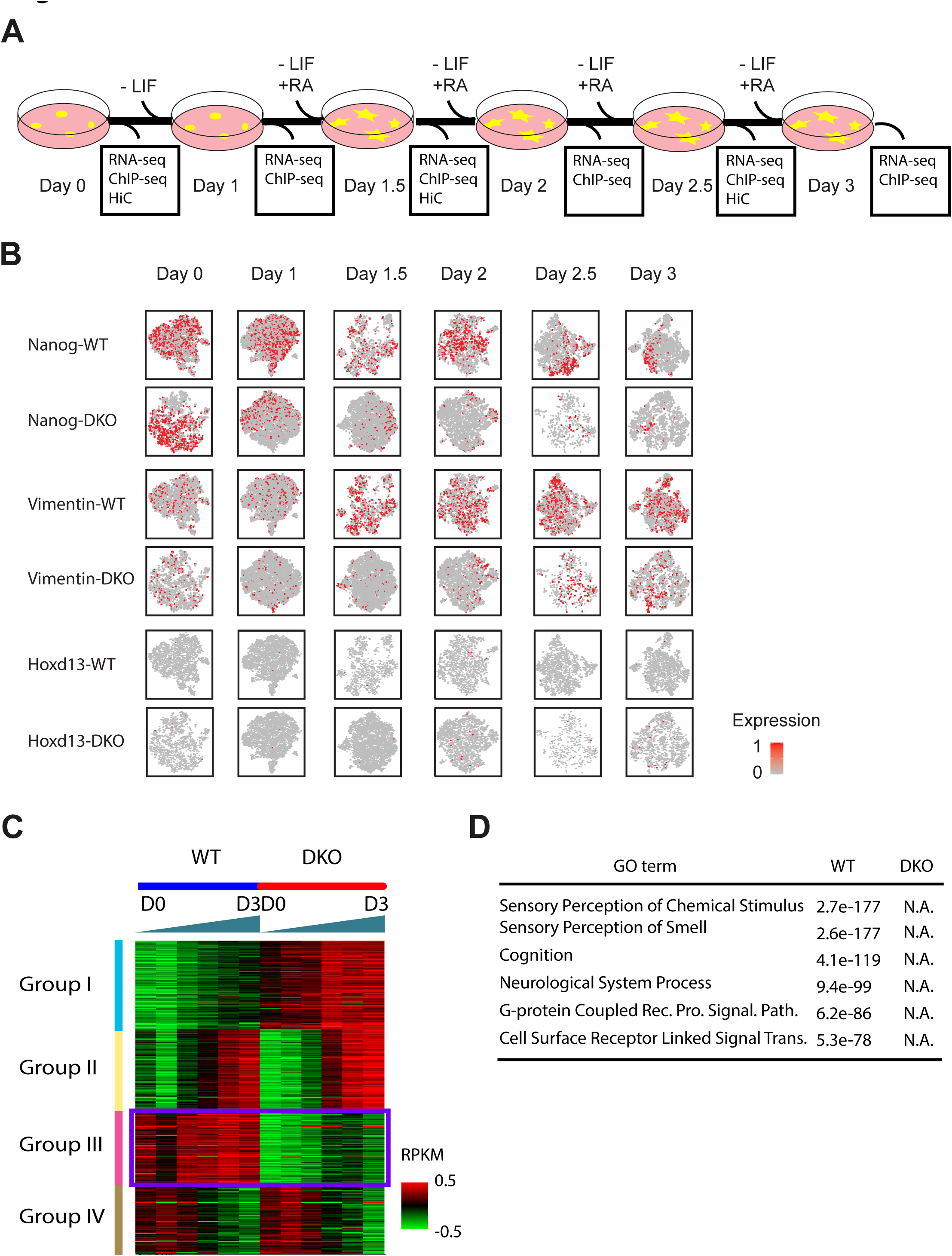
Characterization of gene expression during NPC Differentiation in WT and DKOmESC. **(A)** Scheme of differentiation protocol. Data was collected for RNA-seq and ChIP-seq at every 12-hour time points and Hi-C at every 24-hour time points. **(B)** Single cell RNA analysis of NPC differentiation. Each panel represents 2-D t-SNE (Stochastic Neighbor Embedding) projection of the cells colored by the total UMI counts per cell. x-axis represents t-SNE-1 and y-axis shows t-SNE-2. Red color indicates high expression of the gene that is noted on the left of each row. Each column represents a time point indicated by **(A)**. Expression patterns of Nanog and Vimentin showed that the cell population is well synchronized and no overt subgroup was observed. Nanog, pluripotency marker in embryonic stem cells. Hoxd13 had slight increase in expression in DKO cells. Vimentin, marker for neural progenitor. Hoxd13, posterior HOX transcription factor normally expressed in the posterior part of the body plan during development. **(C)** Clustering analysis of bulk RNA-seq from WT and DKO cells showing that genes could be clustered to 4 different groups depending on the panel of change along differentiation. Note that Group I and Group III genes are mostly MLL3/4-dependent in that they behave differentially between WT and DKO cells. Group II and Group IV are MLL3/4-independent genes and they are expressed in the same pattern in two cell types. **(D)** Gene ontology analysis showing that genes induced only in WT cells are mostly related to neuron function. Note that no GO terms could be detected in genes that are induced only in DKO cells. See also **Table S7**

We also examined the dynamic H3K27ac and H3K4me1 profiles in WT and DKO cells at various time points during NPC differentiation using ChIP-seq (Figure 5A and **Table S3**). Strikingly, at a majority of the distal H3K27ac peaks, H3K4me1 signals were depleted in DKO cells, and cells resulted in a failure to induce cell type specific gene expression. This observation provides further support for a role for MLL3/4 in regulating the chromatin epigenetic landscape at distal enhancers.

**Figure 5.**
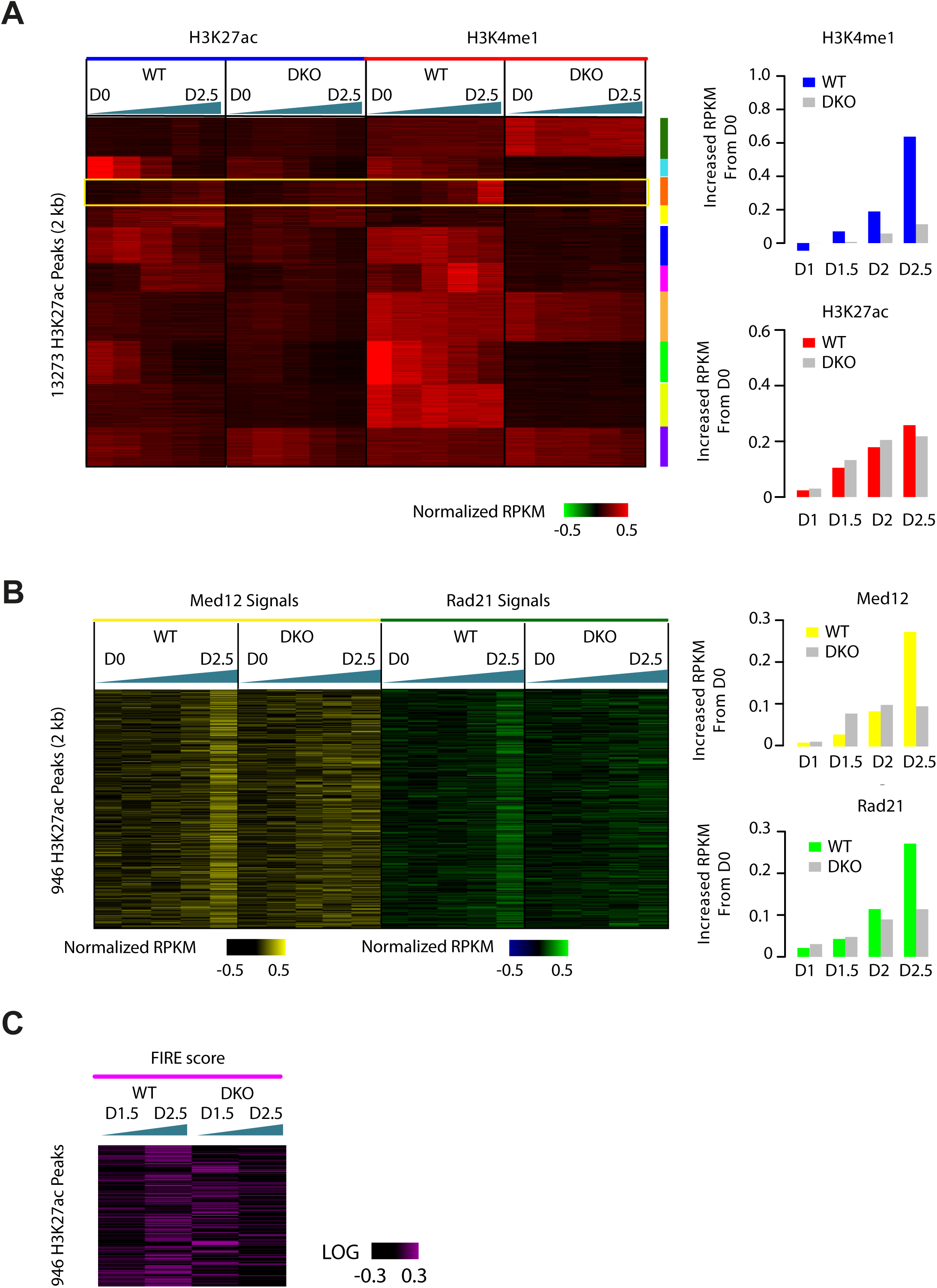
Characterization of H3K4me1, Cohesin and Chromatin Interactions during NPC Differentiation of WT and DKO mESC. **(A)** Clustering analysis showing that H3K27ac peaks could be classified to 10 different groups per change in both H3K27ac and H3K4me1 signal in WT and DKO cells during differentiation. The color key shows the log2 transformed input normalized RPKM value of each peak. Green triangle shows the differentiation time, the thicker side representing later time points. Yellow box emphasizes the group that shows induced H3K27ac and H3K4me1 signal in WT cells along differentiation. The right bar charts show the averaged signal of H3K4me1 (top) and H3K27ac (bottom) of group III. D0, Day1; D2.5, Day2.5. **(B)** Heatmap showing the change of input normalized Mediator subunit Med12 (left) and Cohesin subunit Rad21 RPKM (right) at Group III H3K4me1 peaks emphasized in **(A)**. Note that Cohesin and Mediator binding also tends to increase in WT cells but not in DKO cells. The right bar charts show the averaged signal of H3K4me1 (top) and H3K27ac (bottom) of group III. **(C)** Heatmap showing the change of FIRE score at Group III H3K4me1 peaks emphasized in **(A)**. Note that FIRE score tends to increase in WT cells but not in DKO cells. See also Table S3

To dissect the temporal relationships between H3K4me1 deposition, Cohesin occupancy and chromatin organization, we examined Cohesin and Mediator occupancy and FIRE scores in the regions where accumulating H3K4me1 and H3K27ac signals were detected along differentiation in WT cells but not in DKO **(Figure 5A)**. If H3K4me1 is necessary for Cohesin and Mediator recruitment, gradual loss of H3K4me1 at enhancers during mESC differentiation would lead to a decrease in Cohesin and Mediator binding. As expected, these loci showed increased Cohesin and Mediator binding during differentiation only in WT cells, along with increased H3K4me1 signals (Figure 5B). Interestingly, the FIRE score of this region was also gradually elevated in WT cells, in accordance with the role of MLL3/4 in regulation of chromatin interactions (Figure 5C).

In order to more clearly reveal the concordant change of H3K4me1 signal, Cohesin binding and FIRE score, we zoomed in our analysis to one of our representative locus the *Sox2* SE. We observed a gradual decrease in Cohesin and Mediator binding along differentiation, concurrently with decreased H3K4me1 signals (Figure 6A). Expectedly, the FIRE score spanning this region also gradually decreased, in accordance with the role of H3K4me1 in regulation of chromatin interactions mediated by Cohesin and Mediator (Figure 6B). At another representative locus, we observed accumulation of H3K4me1 but not H3K27ac signals at Day 1.5, while Cohesin and Mediator binding signals were detected at Day 2, 12 hours later than the histone marks (Figure 6C). As expected, FIRE score of this region increased in WT cells, consistent with increased Cohesin and Mediator occupancy (Figure 6D). In DKO cells, neither Cohesin nor Mediator could be detected in both loci at any time point, likely due to lack of H3K4me1 at enhancers. This striking example demonstrated that changes of Mediator and Cohesin binding preceded or coincided with H3K4me1, suggesting that the histone modification could potentially stabilize Cohesin and Mediator loading at enhancers, as well as chromatin organization.

**Figure 6.**
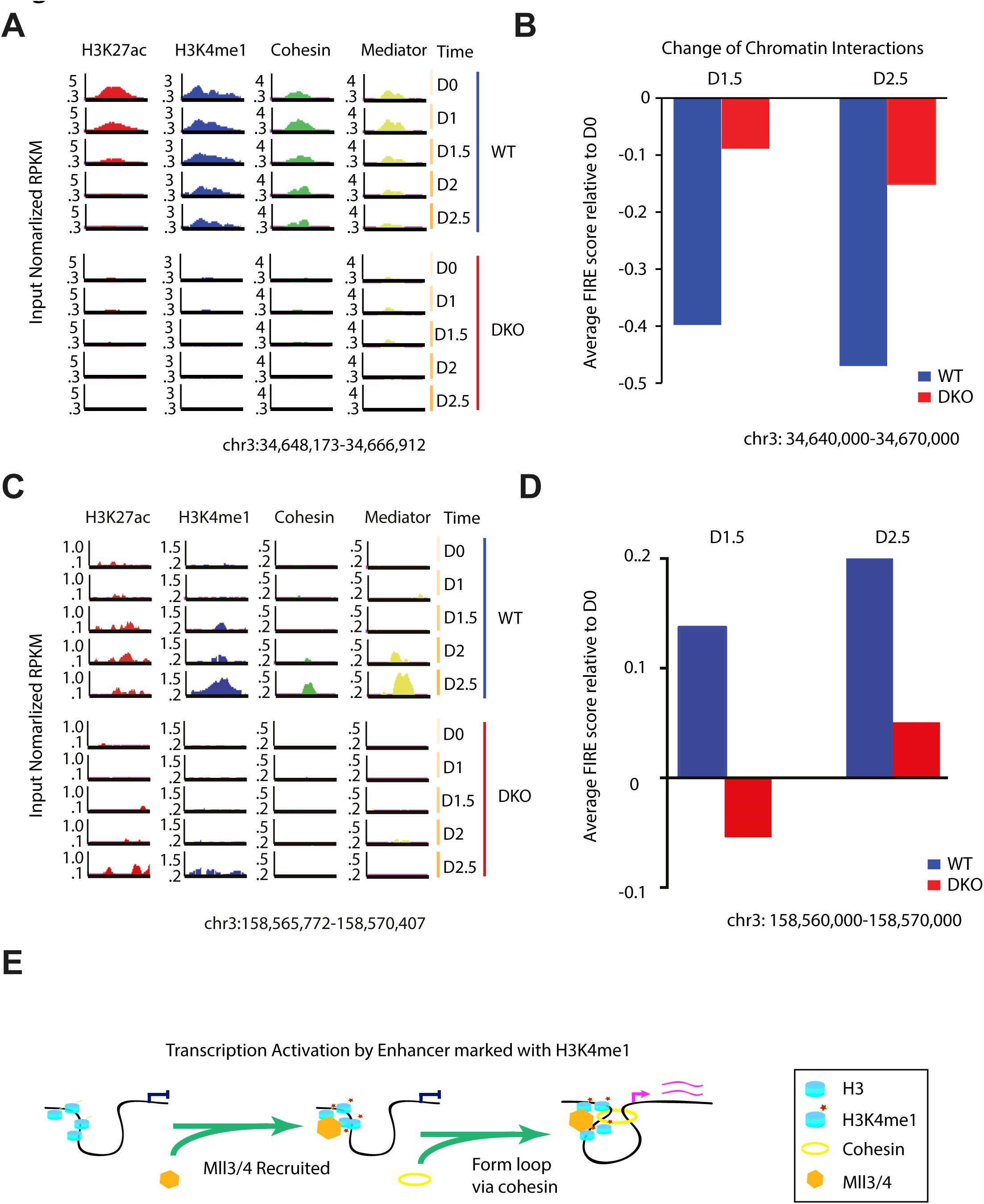
Dynamic histone modification, Mediator and Cohesin Occupancy and Changes in Chromatin Interactions at Representative Loci. **(A)** Genome browser track showing that at *Sox2* SE locus, H3K27ac and H3K4me1 signals were enriched in WT cells but not in DKO cells on Day 0. Along cellular differentiation, H3K27ac and H3K4me1 signals gradually decreased, while Mediator and Cohesin binding were also alleviated. Note that Cohesin and Mediator started decreasing at Day 1.5 while H3K27ac and H3K4me1 both start decreased at Day 1. By contrast, none of the factors could be detected in DKO. Y-axis shows input normalized ChIP-seq reads per kilobase pair per million total reads (RPKM) **(B)** Bar chart showing change of chromatin interactions (FIRE score). Blue bars indicate FIRE score at each time point relative to Day 0. The 30-kb regions are enclosing the entire Sox2 super enhancer in **(A)**. The change of FIRE score in DKO (red bars) was mild compared to WT. **(C)** Genome browser track showing at a representative locus, H3K27ac and H3K4me1 signals gradually increased along cellular differentiation, starting from Day 1.5. Meanwhile, Mediator and Cohesin are also detected to be initiated at Day 2. By contrast in DKO cells, none of the factors could be detected until Day 2.5. Y-axis shows input normalized ChIP-seq reads per kilobase pair per million total reads (RPKM) **(D)** Bar chart showing change of chromatin interactions enclosed in **(C)** (FIRE score). Blue bars indicate FIRE score at each time point relative to Day 0. The 10-kb regions are enclosing the entire 5-kb region shown in panel (A). Less change in FIRE score was observed in DKO (red bars) than in WT. **(E)** A model depicting MLL3/4 and H3K4me1’s role in establishment of chromatin interactions at enhancers. When H3K4 is not methylated at distal enhancer, the promoter is physically separately in different nuclear territory (left). MLL3/4 are recruited to distal enhancers, leading to monmethylation of lysine 4 of histone H3 (middle). Subsequently, the Cohesin complex is recruited to mediate the looping interaction between gene promoter and enhancer. See also Figure S6 and Table S4

### Dynamic Chromatin Contacts at Super-enhancers during ES cell Differentiation Depend on MLL3/4

To more clearly demonstrate the role of MLL3/4 in mediating chromatin interactions, we focused on super-enhancer, which are regions with highly clustered sites of transcription factor and co-factor binding sites, and involved in activing transcription of cell identity genes through long-range chromatin interactions (Dowen et al., 2014). We identified super-enhancers in WT and DKO cells at different time points, based on unusually high levels of H3K27ac signals (Hnisz et al., 2013; Loven et al., 2013; Whyte et al., 2013) (**Supplemental** Figure S6A and **Table S4**). Consistent with a recent report (Schmitt et al., 2016), over half of the super-enhancers identified above were within FIREs in WT mESC, and the H3K27ac signals at these SEs were gradually lost during differentiation. In the DKO cells, the FIRE scores at these regions were near background levels, providing strong support for a role of MLL3/4 in chromatin interactions at FIREs and SEs (**Supplemental** Figure S6B). For example, at the *Sox2* locus, the super-enhancer gradually lost H3K27ac signal until fully depleted by Day 1.5 (Supplemental Figure S6C). FIRE score at this enhancer also decreased along differentiation (**Supplemental** Figure S6D). In DKO cells, the super-enhancer was not properly formed due to lack of MLL3/4 and the chromatin interaction frequency was also very low at this region.

## DISCUSSION

Great strides have been made in the identification of *cis* regulatory sequences in the human genome (Consortium, 2012; Kundaje et al., 2015). Since the majority of the candidate cis-regulatory sequences located far from the transcription start sites, it is generally believed that 3D chromatin architecture plays a critical role in enhancer function. However, the mechanisms by which long-range chromatin interactions are established at lineage-specific enhancers during development are still incompletely understood (Dixon et al., 2015; Phillips-Cremins et al., 2013). In particular, it has yet to be shown whether chromatin remodeling complexes play an active role in chromatin organization at enhancers, despite the data showing close correlation between dynamic histone modifications and chromatin organization during human ES cell differentiation (Dixon et al., 2015; Dixon et al., 2012; Jin et al., 2013; Sanyal et al., 2012; Sexton et al., 2012). Here, we present multiple lines of evidence for a role for histone methyltransferases MLL3/4 in promoting long-range chromatin interactions at enhancers. First, knockout of MLL3/4 at *Sox2* super-enhancer resulted in loss of chromatin interactions between the enhancer and *Sox2* promoter; second, loss of MLL3/4 also led to a decrease in genome-wide chromatin interactions at promoter-distal regions bearing MLL3/4-dependent H3K4me1; third, loss of MLL3/4 resulted in reduced occupancy by the Cohesin complex, which promotes enhancer/promoter interactions in mammalian cells (Kagey et al., 2010). Furthermore, MLL3/4 are required for depositing H3K4me1 and establishing local chromatin interactions at distal enhancers during mouse ES cell differentiation. Finally, pull-down experiment using nuclear extracts showed that the Cohesin complex binds more strongly to H3K4me1-modified nucleosomes than other forms of nucleosomes, and targeted deposition of H3K4me1 in the nucleus enhances recruitment of Cohesin complex to local chromatin. Taken together, our results support a model that MLL3/4 modulate local chromatin interactions at enhancers by depositing H3K4me1 mark and facilitating the binding of Cohesin (Figure 6E).

Cohesin has been shown to mediate chromatin interactions in metazoan cells (Hadjur et al., 2009; Ing-Simmons et al., 2015; Kagey et al., 2010; Mizuguchi et al., 2014; Zuin et al., 2014). Our results suggest that MLL3/4 could facilitate or stabilize Cohesin recruitment at enhancers to promote DNA interactions with other chromatin regions, particularly at promoters. We observed that Cohesin would not bind to the super-enhancers at *Sox2* and *Car2* loci and thus no chromatin interactions formed in the absence of MLL3/4 (Figure 1 and Supplemental Figure S1). During NPC differentiation, we also observed that H3K4me1 signals coincide or precedes Cohesin loading. We provided evidence that the Cohesin may directly or indirectly associate the H3K4me1 mononucleosome. However, we do not yet know how H3K4me1 is recognized by the Cohesin complex. None of the subunits of Cohesin contains known H3K4me1 binding domains. It is likely that a bridging factor might be responsible for Cohesin recruitment to H3K4me1. The ATP-dependent chromatin-remodeling factor SNF2h/SMARCA5 is known to be involved in gene regulation and directly interact with Cohesin component RAD21 (Hakimi et al., 2002). Its interacting factors, ACF/BAZ1A and WSTF/BAZ1B, are known to independently form ACF and WICH complexes with SNF2h/SMARCA5 (Bochar et al., 2000; Ito et al., 1999). BAZ1A/B are known to contain PHD domains, capable of binding to H3 histones, posing them intriguing candidates for H3K4me1 readers (Li et al., 2016). Indeed, we observed that BAZ1A and SNF2h are enriched at Mll3/4-dependent H3K4me1 peaks **(Supplemental Figure S5D)**. Future experiments would be needed to test whether ACF and WICH complexes may play a role in recruitment of Cohesin to H3K4me1 marked nucleosomes. In summary, our model suggests that the enhancer-associated histone modification H3K4me1 is required for enhancer function, either through recruitment of structural complexes that facilitate chromatin interactions or stabilizing the loading of such complexes.

## EXPERIMENTAL PROCEDURES

### Cell Culture

Mouse embryonic stem cell lines were derived from E14 strain and reported separately (Wang et al., 2016a). WT and DKO cells were cultured in mouse ES cell media: DMEM 85%, 15% fetal bovine serum (Hyclone), penicillin/streptomycin, 1× non-essential amino acids (Gibco), 1× GlutaMax, 1000 U/ml LIF (Millipore), 0.4 mM β-mercaptoethanol. Mouse ES cells were initially cultured on 0.1% gelatin-coated petri-dish with CF-1 irradiated mouse embryonic fibroblasts (GlobalStem) and were passaged twice on 0.1% gelatin-coated feeder-free plates before harvesting. Lenti-X 293 cells (Clontech) were cultured in DMEM containing 10% Tet-approved fetal bovine serum (Clontech), penicillin/streptomycin and GlutaMax (Gibco). Alkaline phosphatase staining was performed using the Alkaline Phosphatase Staining kit (STEMGENT) in the presence of MEF feeder cells.

### *in situ* Hi-C

The in situ Hi-C experiments were conducted according to Rao et al., 2014. Briefly, 2 million cells were cross-linked with 1% formaldehyde for 10min at RT and reaction was quenched using 125 mM of Glycine for 5 min at RT. Nuclei were isolated and directly applied for digestion using 4 cutter restriction enzyme MboI (NEB) at 37 °C o/n. The single strand overhang was filled with biotinylated-14-ATP (Life Tech.) using Klenow DNA polymerase (NEB). Different from tradition Hi-C, with *in situ* protocol the ligation was performed when the nuclear membrane was still intact. DNA was ligated for 4 hours at 16 °C using T4 ligase (NEB). Protein was degraded by proteinase K (NEB) treatment at 55 °C for 30 min. The crosslinking was reversed with 500 mM of NaCl and heat at 68 °C o/n. DNA was purified and sonicated to 300-700 bp small fragments. Biotinylated DNA was selected with Dynabeads MyOne T1 Streptavidin beads (Life Tech.). Sequencing library was prepared on beads and intensive wash was performed between different reactions. Libraries were checked with Agilent Bioanalyzer 2100 and quantified using Qubit (Life Tech.). Libraries were sequenced with Illumina Hiseq 2500 or Hiseq 4000 with 50 or 100 cycles of paired-end reads.

### Data Processing and FIRE Analysis

Hi-C pre-processing pipeline is applied to all the raw Hi-C data as described in (Dixon et al., 2015). All intra-chromosome reads within 15-kb are removed from the downstream analyses. The filtered intra-chromosome reads connecting two genomic regions located greater than 15-kb were selected and binned into 10-kb resolution to build the Hi-C contact matrices. For each 10-kb bin, the raw FIRE score is defined as the total number of chromatin interactions within 200-kb. A custom normalization pipeline, ‘HiCNormCis’, was modified from the HiCNorm (Hu et al., 2012) and applied to normalize raw FIRE score. Briefly, a Poisson regression model is fitted for each 10-kb bin, with the raw FIRE score as the outcome variable, and three local genomic features, such as effective fragment length, GC content and mappability score, as the covariates. The residuals from the Poisson regression model are used as the normalized FIRE score, which are comparable among different 10-Kb bins and different Hi-C datasets. Next, for each 10-kb bin, z-score is calculated based on the normalized FIRE score. 10-kb bins with z-score > 1.64 (p-value < 0.05) are defined as FIRE bins. More details could be found in **Extended Experimental Procedure**.

### 3D-FISH

WT or DKO cells were cultured on laminin-coated coverslips (Neuvitro) for 1 hour at 37°C, and then rinsed with PBS and fixed with 4% PFA in PBS for 10 minutes. The fixation was quenched with 0.1 Tris-HCl, pH 7.5, for 10 minutes, rinsed and stored in PBS.

To generate probes, fosmid clone spanning Sox2 SE locus or promoter locus (BACPAC), was labeled with Alexa-fluor 568-5-dUTP or Alexa-fluor 488-5 dUTP (Life Technologies) using Nick Translation Kit (Roche) and incubated at 15°C for 4 hours. The reaction was stopped by 1 µl 0.5M EDTA, pH 8 and heat-inactivated at 65°C for 10 minutes. Unbound dyes were removed using illustra ProbeQuant G-50 Micro Columns (GE Healthcare) following the manufacturer’s instructions. For each hybridization, 20 ng of FISH probes were ethanol precipitated with 10 µg of sonicated salmon sperm DNA, 4 µg of mouse Cot-1 DNA, 1/10th volume of 3M sodium acetate, pH 5.2, and 2.5 volume of 100% ethanol. Each probe was precipitated and dissolved in 5 µl of formamide and 5 µl of 2X hybridization mix (8X SSC/40% dextran sulfate) at 55°C for 20 minutes. The probes were denatured at 75°C for 5 minutes before applying to slides.

Fixed cells on coverslips were blocked in 5% BSA/ 0.1% Triton-X/1X PBS for 30 minutes at 37°C and washed with 0.1% Triton-X 100 in PBS. Cover slips were then permeabilized in 0.1% saponin/0.1% Triton-X/1X PBS for 10 min at room temperature, incubated in 20% glycerol in PBS for 20 minutes at room temperature, freeze-thawed three times in liquid nitrogen, incubated in 0.1M HCl for 30 minutes at room temperature, blocked in 3% BSA and 100 µg/ml RNase A in PBS for 1h at 37°C, permeabilized again in 0.5% saponin/0.5% Triton-X 100/1X PBS for 30 minutes at 37°C, and washed in 2X SSC. Cells were denatured with 70% formamide/2X SSC at 73°C for 2.5 minutes and with 50% formamide/2X SSC at 73°C for 1 minute, after which denatured probes was applied to the slide. After overnight incubation at 37°C, cells were washed twice with 50% formamide/2X SSC at 37°C for 15 minutes and 2X SSC at 37°C. The cover slips were stained with DAPI and rinsed in PBS before mounting on the slides with ProLong Gold Antifade Mountant and sealed with nail polish. Images were acquired at 100x magnification on DeltaVision RT Deconvolution microscope, controlled by SoftWorX software. DNA spots were identified and measured using TANGO v.0.93 software.

### RNA-seq, single cell RNA-seq and Data Analysis

Total RNA from ES cells was extracted with Trizol^®^ according to protocol (Thermo Scientific, 15596-026). PolyA+ RNA was purified with the Dynabeads mRNA purification kit (Life Tech.). The mRNA libraries were prepared for strand-specific sequencing using Illumina TruSeq Stranded mRNA Library Prep Kit Set A (Illumina, RS-122-2101) or Set B (Illumina, RS-122-2102). Libraries were sequenced with Illumina Hiseq 2500 for 100 cycles single reads.

For single cell RNA-seq, cells were harvested after trypsin treatment and washed with PBS. For each time point, cell densities were estimated using a hemocytometer and 2000-5000 cells were collected for library preparation. The single cell library was prepared with Chromium™ Single Cell 3’ v2 Library kit (10XGenomics). The libraries were sequenced using Illumina Hiseq4000 and 16-20 million single read reads were acquired for each individual library respectively. The single cell RNA-seq data was analyzed using Cellranger R kit that was provided by 10XGenomics with default parameters. The quality control statistics was listed in **Supplemental Table S7**.

Sequence reads were mapped to mouse mm9 reference genome with TopHat (Trapnell et al., 2009). The differential expression was analyzed with Cuffdiff (Trapnell et al., 2012). We plotted the differentially expressed genes if the fold change of adjusted FPKM value between E14 and DKO is larger than 2.

Gene Ontology analysis was carried out using DAVID release 6.7 with default parameters (Huang da et al., 2009).

Additional experimental information can be found in “**Supplemental Information**”.

## EXTENDED EXPERIMENTAL PROCEDURE

### 4C-seq and Data Analysis

4C-seq experiments and analysis was performed as described previously(van de Werken et al., 2012a). Briefly, 5 million cells were cross-linked with 2% formaldehyde for 10 min at room temperature (RT) and quenched by adding 125 mM Glycine with 5 min additional incubation at RT. Cells were lysed and nuclei were isolated and digested with Csp6I (Thermo Scientific) or Dpn II (NEB) over night (o/n). Enzyme was inactivated by heat at 65 °C for 20 min. The digested chromatin was subjected for ligation for 16 h with T4 ligase (Life Technologies). DNA was then purified with phenol/chloroform extraction and ethanol precipitation before the second digestion with NlaIII (NEB) or BfaI (NEB) at 37 °C o/n. After enzyme inactivation, a second ligation was performed at 16 °C for 4 h and DNA was purified, of which 4.8 µg in total was used for PCR amplification using 4 different pairs of primers (**Table S5**) which were designed compatible for illumina Hiseq 2500 sequencer.

Sequencing data were analyzed using a custom pipeline ‘4Cseq’ as previously described(van de Werken et al., 2012b) with all default parameters. For consistency, all sequencing data involving E14 and DKO genomes in this work were mapped to mouse mm9 reference genome. Contact frequency was visualized for genomic regions in a 300 kb window that includes both SOX2 gene and SOX2 SE (chr3: 34,448,927-34,765,152).

### Lentiviral Packaging and shRNA Knock-Down

Lentiviral particles were prepared using the Lenti-X single shot packaging system (Clontech), according to the manufacture’s guidelines. Control shGFP (Addgene #30323) and murine shRad21 (Sigma SHCLNV_NM009009 (TRCN0000176084)) plasmids (Target sequences see **Table S5**) were transfected to Lenti-X 293 cell line (Clontech) and viral supernatant were concentrated using the Amicon Ultra-15 100 kDa centrifugal filters (Millipore). For shRNA knockdown, E14 cells were plated on a 6-well dish coated with 0.1% gelatin the day before transduction (2×10^5^ cells per well). Concentrated viral supernatant were added to mouse ES medium containing 8 µg/mL polybrene (Sigma). Media containing lentiviruses were replaced with fresh media 24 h post-infection. Infected cells were selected by puromycin (1 µg/mL) for 72 h.

### Chromatin Immunoprecipitation Followed by Sequencing (ChIP-seq) and Data Analysis

The ChIP-seq has been carried out as previously described (Jolma et al., 2013; Tuupanen et al., 2012; Yan et al., 2013). Briefly, 2 million cells were crosslinked with 1% formadehyde for 10 min at RT. The reaction was quenched by adding 125 mM of Glycine and incubating for 5 min at RT. Cells were lysed in RIPA buffer (10 mM Tris-HCl pH 8.0, 140 mM NaCl, 1 mM EDTA, 1% Triton X-100, 0.1% SDS, 0.1% sodium deoxycholate) supplemented with protease inhibitor (Roche). And chromatin was sonicated into short fragments (300-700 bp). The fragmented chromatin was incubated with antibodies (**Table S6**) to pull down the specific DNA bound TFs or histones. After intensive wash, DNA was purified and prepared as sequencing library using illumina Truseq LT kit. Several samples with different indexes were pooled together for 50 or 100 cycles single read sequencing with illumina Solexa sequencer or Hiseq 2500.

Sequencing reads were mapped to mouse mm9 reference genome using bowtie (Langmead et al., 2009). PCR duplicates were removed and peaks were called with MACS (Zhang et al., 2008) using input chromatin as control, with the parameter-m 5, 50 otherwise default. RPKM was calculated for each peak by dividing the number of reads overlapping peak with the length of the peak and multiply 1000, and the resulted value would be multiplied with a scale factor as to normalize the total number of reads to 1 million. The RPKM of each peak for samples was subtracted with the RPKM of that peak for input. If the subtracted number is less than 0, the RPKM of the peak for that sample will be assigned as 0.

In order to define Mll3/4 dependent H3K4me1 peaks, we merged H3K4me1 peaks from both E14 and DKO cells and extended all peaks to 2 kb wide. We merged two close peaks if they were partially overlapping. According to the RPKM described above, we calculated RPKM for all the 2-kb peaks for both E14 and DKO cells. We sorted the peaks according to the difference of the RPKM between E14 and DKO. If the RPKM of a given peak in E14 is over 0.6 larger than DKO, that peak will be classified to ‘Decreased’. Similarly, if the difference is over 0.6 smaller than DKO, that peak will be classified as ‘Increased’. Other peaks will be classified as ‘Non-differential’.

The sequences from the top 300 decreased peaks were submitted to AME (MEME-suite (Bailey et al., 2009)), using the bottom 300 increased peaks as background.

GO analysis for different categories of H3K4me1 peaks was performed with GREAT (McLean et al., 2010).

### RNA-seq and Data Analysis

Total RNA from ES cells was extracted with Trizol^®^ according to protocol (Thermo Scientific, 15596-026). PolyA+ RNA was purified with the Dynabeads mRNA purification kit (Life Tech.). The mRNA libraries were prepared for strand-specific sequencing using illumina TruSeq Stranded mRNA Library Prep Kit Set A (illumina, RS-122-2101) or Set B (illumina, RS-122-2102). Libraries were sequenced with illumina Hiseq 2500 for 100 cycles single reads.

The single cell RNA-seq was carried out using Chromium™ Single Cell 3’ v2 Library (10XGenomics). For each time point, 2000-5000 cells were analyzed.

Sequencing was mapped to mouse mm9 reference genome with Tophat (Trapnell et al., 2009). The differential expression was analyzed with Cuffdiff (Trapnell et al., 2012). We plotted the differentially expressed genes if the fold change of adjusted fpkm value between E14 and DKO is larger than 2.

Gene Ontology analysis was carried out using DAVID release 6.7 with default parameters (Huang da et al., 2009).

### RNA Extraction and qPCR

Total RNA was isolated from harvested cells using the RNeasy columns (Qiagen) according to the manufacturers instructions. cDNA were synthesized from 400 ng of total RNA using High Capacity cDNA Reverse Transcription kit (Applied Biosystems). qPCR was performed in triplicates using SYBR FAST qPCR master mix (KAPA biosystems) on the LightCycler 480 (Roche). Two independent sets of qPCR primers for mouse Actb and Sox2 were used: customer synthesized primers (**Table S5)** and commercially available primer sets for mouse Actb (Qiagen, catalogue no. PPM02945B) and Sox2 (Qiagen, catalogue no. PPM04762E), and a set of primers for Rad21 (Qiagen, catalogue no. QT00141204).

### Nucleosome Assembly and Pull down assay

Histones are expressed using E.coli strain BL21 (DE3) transformed with cDNA of wild type H2A, H2B, H3, H4 and a mutant H3 C110A K4C contruct (a generous gift from Dr. M. Carey). In order to make methyl-lysine analogs, we used a previously described protocol (Simon et al., 2007). Briefly, 5 mg of H3 was incubated and mixed with (2-hal-oethyl) amines under reducing conditions, followed by being quenched with β-mercaptoethanol. The methylated histone was dialyzed against water overnight, and spun to remove precipitant. Equimolar amounts of histones were mixed under denaturing conditions and dialyzed overnight to assemble octamers followed by size selection (Luger et al., 1999).

Biotin tagged double stranded 601λ positioning DNA sequence was prepared as previously described (Dyer et al., 2004). The mono-nucleosomes were produced via serial salt dialysis (Carruthers et al., 1999). The H3 lysine 4 methylation was tested by western blotting with antibodies specifically recognizing various H3K4me states.

The different modified mono-nucleosomes were immobilized to streptavidin-coated beads (Invitrogen MyOneT1) as per manufacturers instructions and used as baits in following binding studies. Briefly, three micrograms of mono-nucleosomes were pre-bound to MyOneT1 beads. Immobilized nucleossomes were incubated with rotation with HeLa Nuclei Extract (200 µl of ~5 mg/ml) for 1 hour at room temperature. Beads were washed 3 times with wash buffer containing 250mM NaCl, 25mM Tris pH 8.0, 1mM EDTA, 0.2% NP40, and 1mM DTT and resuspended in equal volume of 2X Laemmli Sample Buffer (Bio-Rad). Binding was tested via western blotting using antibodies listed in **Table S6**.

### Hi-C FIRE Score Clustering

We used R function ‘hclust’ with the complete linkage to carry out hierarchical clustering analysis. In specific, we first perform log2 transformation to the FIRE score, and then calculated the Euclidean distance between any two samples.

### Topological Associating Domain and Boundary calling

Topological domains were called based on the directionality index (DI) score using a Hidden Markov Model (HMM) as previously described (Dixon et al., 2012). The software used can be downloaded at Hi-C Domain Caller. According to the domain patterns, the genome is partitioned as follows: domains are marked as domains; gaps between domains that are larger than 100 kb were marked as unstructured regions; gaps between domains that are smaller than 100 kb were marked as boundaries; if two domains are consecutive, the 10 kb window centered at the boundary is marked as a boundary.

### Support Vector Machine for FIRE Classification

We first partitioned the genome into non-overlapping bins of the same length (10kb) and filtered those with poor mappability. Bins with z-score greater than 1.65 (*P* < 0.05) were selected as positive hits, same amount of bins with smallest z-score were chosen as negative set. With 8 features, Support Vector Machine (SVM) implemented by R package ‘1071’ was applied to classify positive FIRE bins from the negative ones with default model setting (gamma=1, epsilon=0.1 and radical kernel), prediction performance was evaluated by AUC (Area Under ROC curve) using 5-fold cross validation. To further evaluate the importance of each variable, we made prediction based on the same setting but using one feature each time. AUC estimated by 5-fold cross validation for each feature reflects its decimation power.

### Neural Progenitor Cell Differentiation

NPC differentiation protocol is adapted from previously published methods with modification (Bibel et al., 2004; Hon et al., 2014; Wang et al., 2012). Briefly, mouse embryonic stem cell line WT and DKO cells were grown on *γ*-irradiated Mouse Embryonic Fibroblast feeder cells before seeding. Cells were split and seeded to 10-cm petri-dish coated with 0.2% Gelatin type-A one day before differentiation in ES culture medium supplemented with LIF. On Day 0, LIF was deprived from the culture medium and cells were continued to be cultured for 24 hours. From Day 1 to Day 3, cells were cultured in LIF-deprived ES medium supplemented with 5 µM retinoic acid (RA). Cells were harvested every 12 hours and aliquoted for further assays. One million cells were collected for RNA-seq, two millions cells were collected for in situ Hi-C experiments, fixed 1% formaldehyde (sigma). Five million cells were collected for 4C-seq and ChIP-seq, fixed with 2% and 1% formaldehyde, respectively.

### Super-enhancer Analysis

Super-enhancer is defined using H3K27ac ChIP-seq data, similar to (Hnisz et al., 2013). Briefly, MACS was used to call narrow peaks of H3K27ac with input as controls. The peak file was then used as a guide file to define Super-enhancer, using published algorithm ROSE (Hnisz et al., 2013; Loven et al., 2013). For each ChIP-seq library including H3K27ac and input at each time points, the number of uniquely mappable reads of 15 million was set as a minimum requirement for super-enhancer call.

### HiCNormCis and FIRE calling

We developed a novel computational approach, named as HiCNormCis (Schmitt et al, manuscript under minor revision at Cell Reports), to remove systematic biases in total cis intra-chromosomal interactions. We first filtered out all intra-chromosomal interactions with 15kb since they are very likely to be self-ligation artifacts. Next, we divided the mouse reference genome (mm9) into 10kb bins, and for each 10kb bin, calculated the total cis intra-chromosomal interactions within 200kb. Consistent with the previous study (Yaffe and Tanay, 2011), we observed that the raw total cis intra-chromosomal interactions contain biases from three local genomic features, including restriction enzyme fragment length, GC content and mappability score. We applied a Poisson regression approach to remove three systematic biases. In specific, let *y_i_* represent the raw total cis intra-chromosomal interactions at the *i* th 10kb bin. In addition, let *F_i_*, *GC_i_* and *M_i_* represent the effective fragment size, GC content and mappability score at the *i* th 10kb bin, respectively. The definition of these three local genomic features is described in our previous work (Hu et al, 2012). Assume *y_i_* follows a Poisson distribution with mean *θ_i_*, we fitted a Poisson regression model 
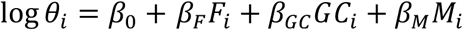
, where *β_F_*, *β_GC_* and *β_M_* are regression coefficients of the effective fragment size, GC content and mappability score respectively. Here *β_0_* is a Poisson offset to account for total sequencing depth. After fitting this Poisson regression model, we obtained the estimate of the unknown parameters 
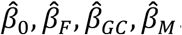
 Next, for each *i* th 10kb bin, we defined residual 
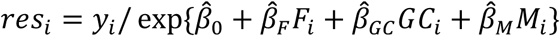
 as the FIRE score. We further converted FIRE score into z-score and corresponding one-sided p-value. 10kb bins with one-sided p-value less than 0.05 are determined as FIRE bins. We have shown that the raw total cis intra-chromosomal interactions *y_i_* show strong correlation with three local genomic features *F_i_*, *GC_i_* and *M_i_* (**Supplemental** Figure S3E). After applying HiCNormCis, the FIRE score *res_i_* show negligible correlation with these genomic features (**Supplemental** Figure S3F), indicating that HiCNormCis has successfully removed such biases. As a comparison, we also implemented two popular matrix balancing based Hi-C data normalization approaches, Vanilla Coverage (Rao et al, 2014) and ICE (Imakaev, et al, 2012). However, both Vanilla Coverage and ICE normalized total cis intra-chromosomal interactions still show high correlation with *F_i_*, *GC_i_* and *M_i_* (**Supplemental** Figure S3G, **S3H**). Therefore, we conclude that VC and ICE are not optimal for correcting biases buried in total cis intra-chromosomal interactions, and decide to use HiCNormCis for Hi-C data normalization.

## Supplemental Figure Legends

**Figure S1.**
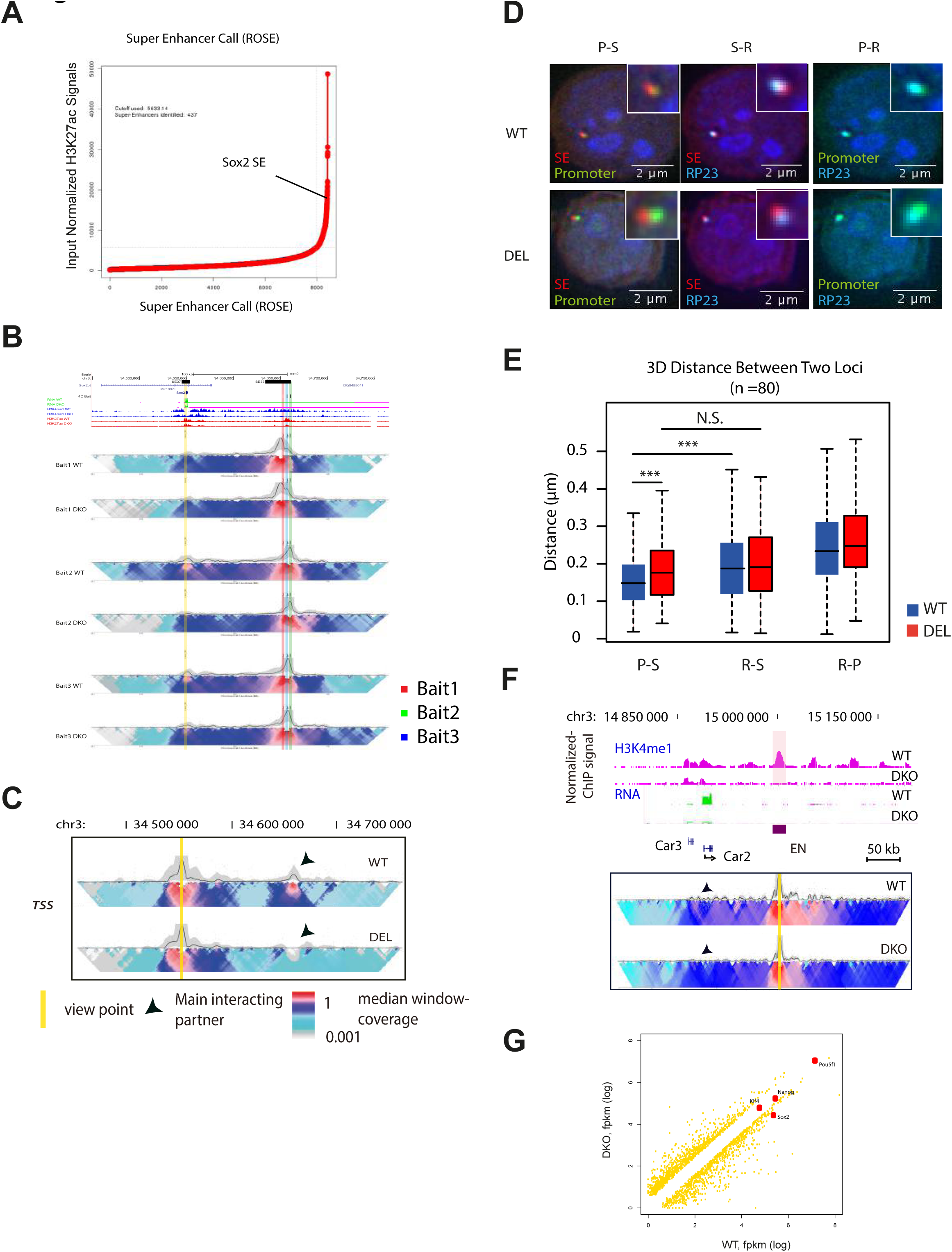
Mll3/4 Deficient mESCs Reveal Lost Interaction between Sox2 Gene Body and SE, Related to Figure 1. **(A)** Super-enhancer call using H3K27ac ChIP-seq data in WT cells. x-axis shows the rank of enhancers, and larger number represents higher rank of peaks with H3K27ac signals. y-axis is the input normalized H3K27ac ChIP-seq coverage within the peak. **(B)** Similar to Fig. 1b, the 4C-seq data from different viewpoints shows that the interaction between *Sox2* gene body and SE is lost upon Mll3/4 depletion in mESCs. **(C)** A 2D-heat map of 4C-seq analysis shows a significant reduction in contact frequency between *Sox2* TSS and *Sox2*-SE in DEL cells compared to F123 WT cells. **(D)** Microscopy images of 3D FISH shows that physical distance between *SOX2* SE and promoter becomes larger in DEL cells than F123 WT. Red dots, probes hybrid to SE locus, Green dots, probes detecting promoter locus; Cyan dots, probes detecting a region (RP23 locus) that is located 170 kb downstream from SE. **(E)** Summary of FISH data from 80 different cells for both cell types respectively. Note that distance between promoter and SE was significantly larger in DKO than WT. In WT cells, SE is significantly closer to promoter than to RP23 while the difference was not observed in DEL cells, indicating the loss of interaction between SE and Promoter in DEL cells. S-R, distance between RP23 and SE; P-R, distance between RP23 and Promoter; P-S, distance between Promoter and SE. Asterisks indicate statistical significance tested with student t-test (*** p<0.005). **(F)** 4C-seq shows that the enhancer interacts with *Car2* gene in WT cells but not in DKO cells (lower). The H3K4me1 ChIP-seq tracks are shown as a reference (upper). Note that H3K4me1 signals at both *Car2* enhancer and *Car2* gene body are decreased in DKO cells. EN indicates the location of the enhancer. **(G)** RNA-seq data shows that except Sox2, transcription level of other pluoripoteny factors (labeled with red dot) is not changed. FPKM is computed using cufflink with three biological replicates.

**Figure S2.**
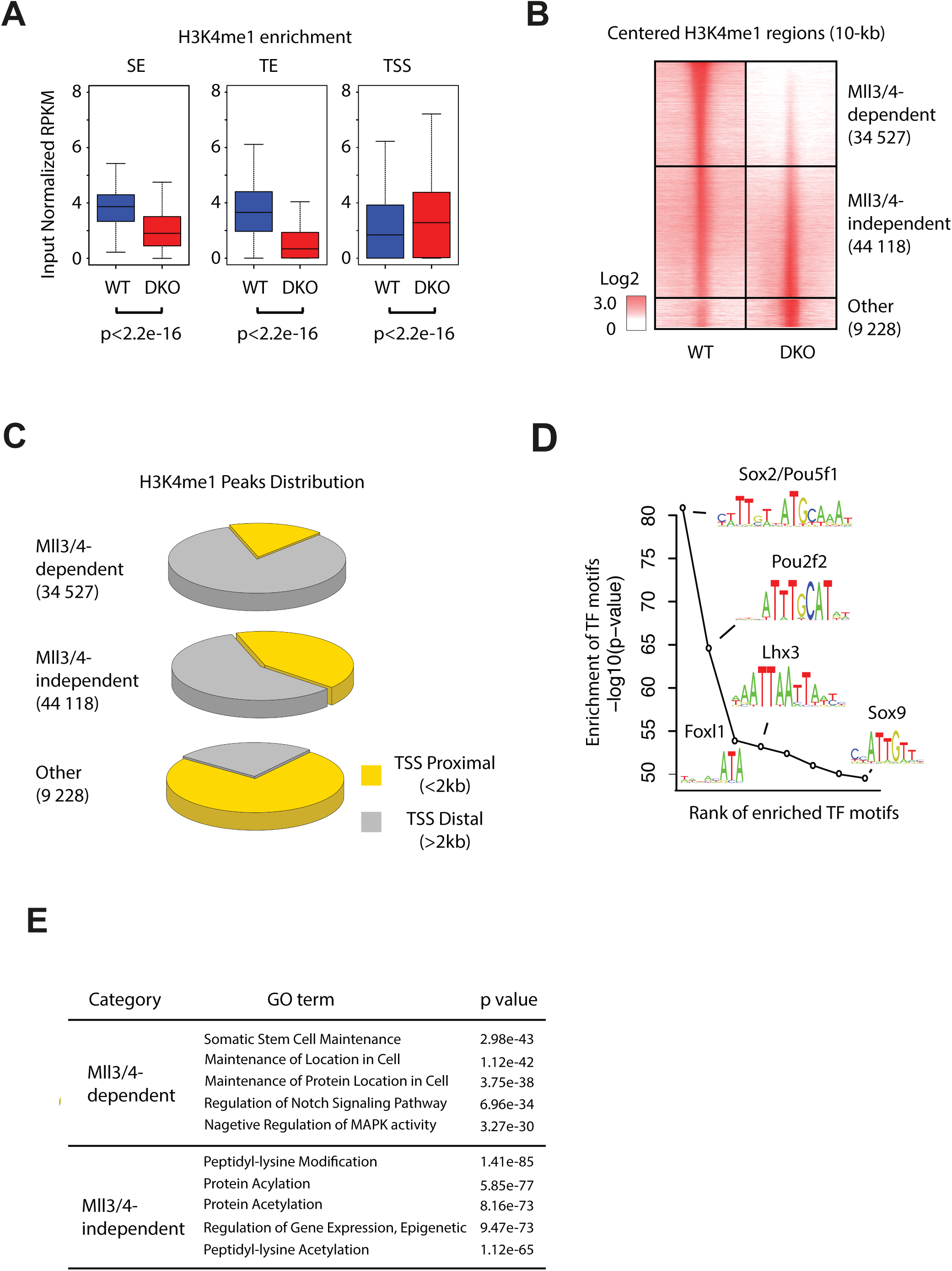
Mll3/4 Are Required for Genome-wide Deposition of H3K4me1 at promoter-distal Enhancers, Related to Figure 1. **(A)** Comparison of H3K4me1 ChIP-seq signals between WT and DKO cells at cis-regulatory elements (TE and SE, defined by (Whyte et al., 2013) and within 2 kb of TSS. SE, super-enhancer. TE, typical enhancer. TSS, transcription starting site. Median values were shown as vertical lines in the box plots, the upper edge and lower edge show the 25th and 75th quantile, and error bars indicate 10th and 90th quantile. p value was computed with two-tail Student t-test. **(B)** A Venn Diagram (left) shows the overlap of H3K4me1 ChIP-seq peaks between WT (blue) and DKO (red) cells. Numbers of Mll3/4-dependent H3K4me1 regions, Mll3/4-independent H3K4me1 peaks and other H3K4me1 regions are shown. Heatmap (right) shows the distribution of H3K4me1 ChIP-seq signal within 10 kb of peak summits in WT and DKO cells respectively. Each row represents the same 10-kb window surrounding the peak summit in E14 and DKO cells. Color key shows the log2 transformed RPKM. **(C)** Distribution of H3K4me1 regions in different genomic positions relative to TSS. **(D)** Result of motif enrichment analysis performed with the top 500 Mll3/4-dependent H3K4m1 peaks as foreground and the bottom 500 Mll3/4-independent H3K4m1 peaks as background. Note that motif for Sox2, a master regulator of embryonic stem cell pluoripotency, has the lowest p value. **(E)** Shown is the result of Gene Ontology analysis using all Mll3/4-dependent H3K4me1 regions as foreground and all H3K4me1 regions as background.

**Figure S3.**
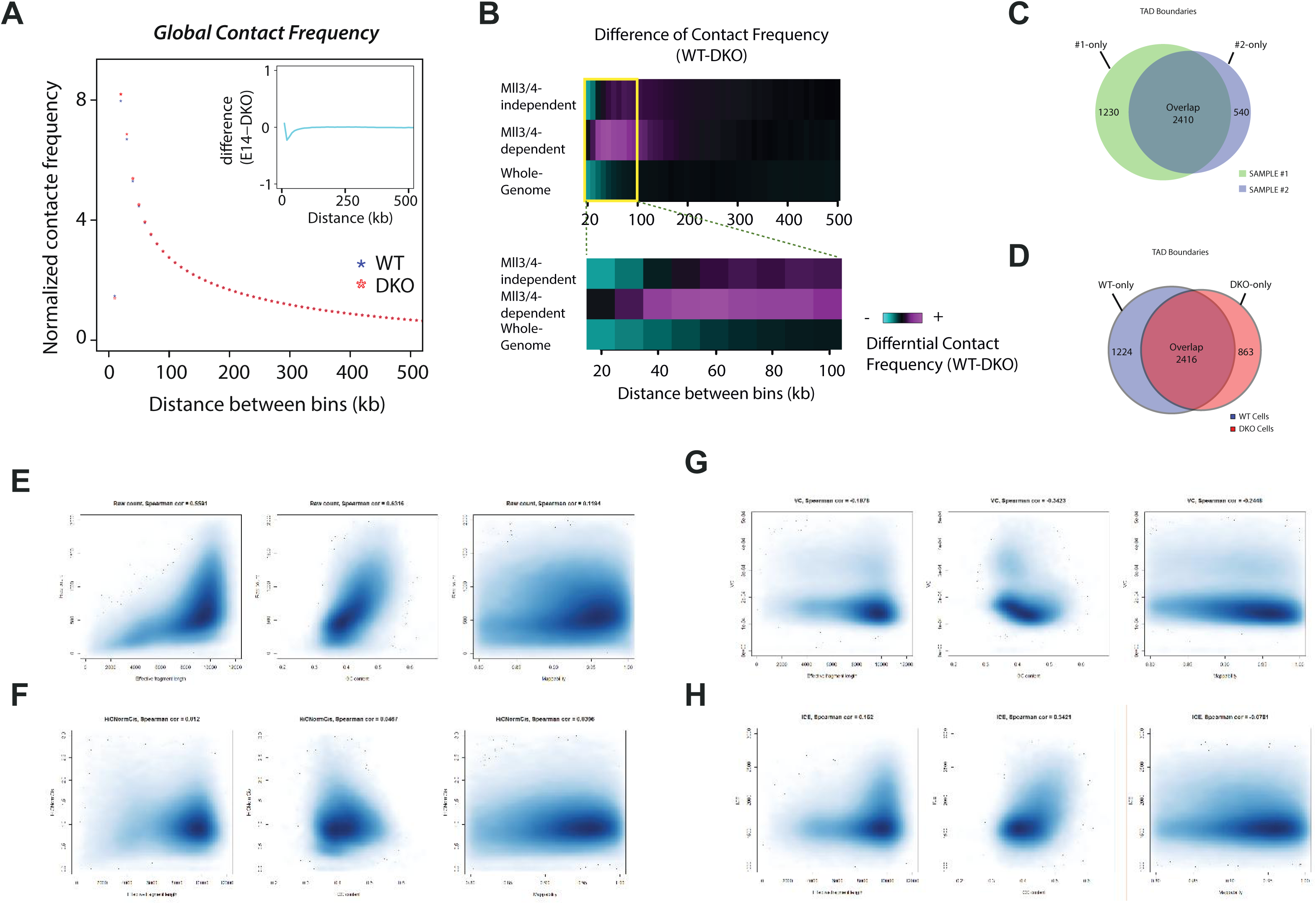
H3K4me1 does not Affect Global Contact Frequency but is Required for FIREs, Related to Figure 2. **(A)** Global contact frequency is not obviously affected by Mll3/4 depletion. x-axis shows the distance between two bins. Y-axis is the average value of normalized Hi-C contact frequency. Blue stars, data from WT cells. Red stars, data from DKO cells. Inset shows the subtracted values between two cell types. **(B)** Heatmap showing the difference in normalized intra-chromosomal contact frequency at various distance between WT and DKO cells for chromatin regions among different classes of H3K4me1 regions. Mll3/4-independent bins include only Mll3/4-independent H3K4me1 regions; Mll3/4-dependent bins include only Mll3/4-dependent H3K4me1 regions; Whole-Genome, all bins for comparison. Color key shows the normalized differential contact frequency. **(C)** Comparison of TAD boundaries between two biological replicates of WT cells. Note that the majority of TAD boundaries are shared by the two samples. **(D)** Comparison of TAD boundaries between WT and DKO cells. Note that the difference is of the similar magnitude as biological replicates shown in panel **(C)**. **(E)** Scatter plot shows that the raw counts of Hi-C libraries are correlated with fragment length (left), GC content (middle) and mappability (right). **(F)** Scatter plot shows after HiCNormCis normalization, the interaction frequency is no longer correlated with the three main bias factors mentioned in **(E)**. **(G)-(H)** Similar to (**F**), Scatter plot shows after VC or ICE normalization, the interaction frequency is still correlated with the three main bias factors mentioned

**Figure S4.**
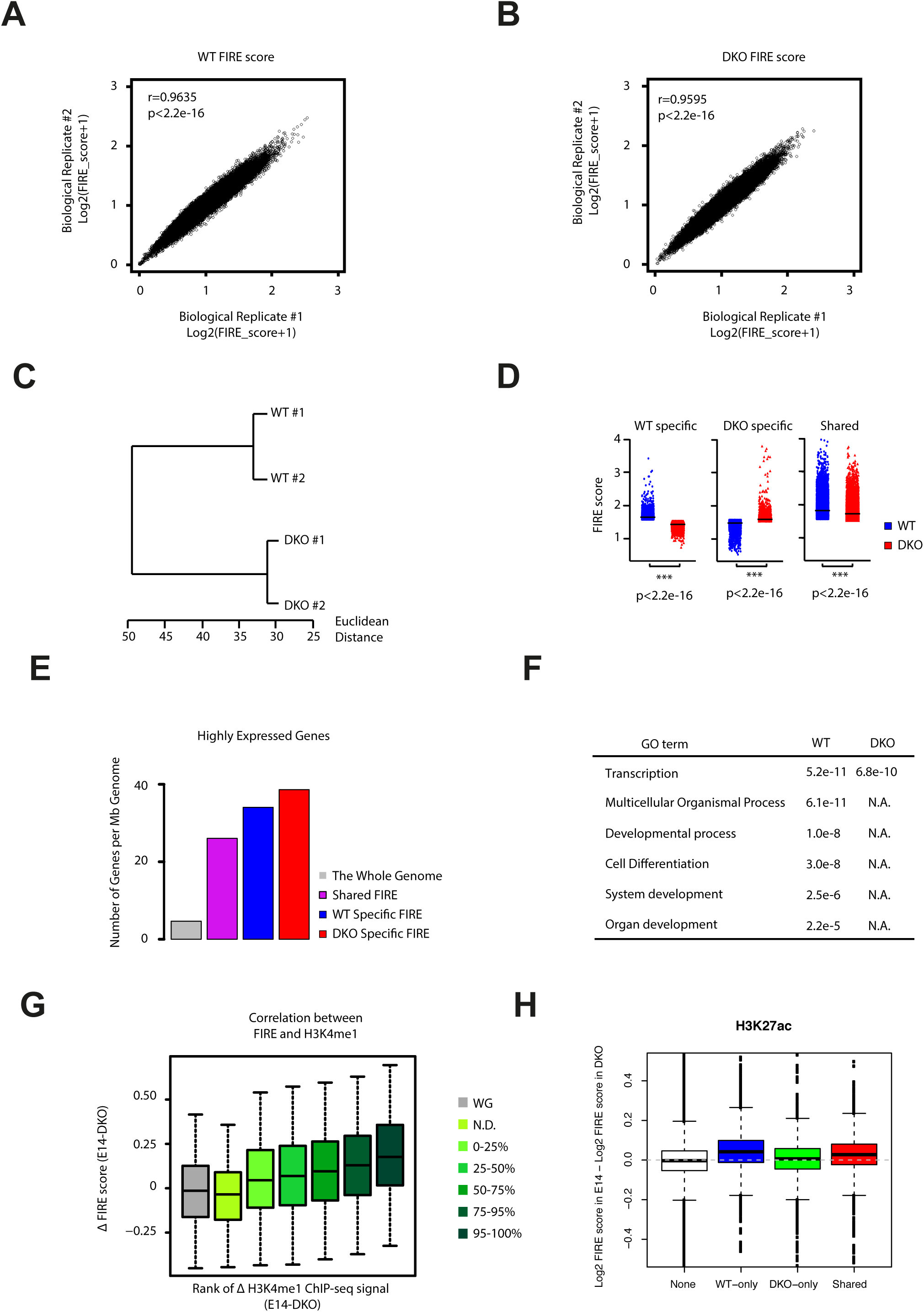
H3K4me1 is Required for FIREs, Related to Figure 2. **(A, B)** Scatter plots show comparison of FIRE score between two biological replicates of WT cells **(B)** and DKO cells **(C)**, respectively. Insets show the Pearson correlation coefficient r and p value. **(C)** Cluster analysis using Pearson correlation coefficient r between samples. The x-axis represents the maximum possible Euclidean distance between samples belonging to two different clusters. From the dendrogram, we observed that two biological replicates of the same condition first clustered together, indicating that the variation between two biological replicates of the same condition is less than the variation between wild type cells and mutant cells. **(D)** Jittered scatter plot shows FIRE score of different categories of FIREs. Note that in the shared FIREs, WT cells show higher FIRE score than DKO cells. The p value is computed with Mann-Whitney U test. **(E)** The gene density of expressed genes (FPKM>=1) per Mb DNA sequences in FIREs are significantly higher than the average genome. **(F)** Gene ontology analysis of genes located in WT and DKO specific FIREs repspectively. The genes in WT-specific FIREs are more functionally related to cell differentiation and development. The Benjamini-adjusted p values are shown in each category. **(G)** Boxplot shows the correlation between the change of H3K4me1 and change of FIRE score throughout the genome. The change of H3K4me1 is classified in different quantiles according to the change of the input normalized RPKM value. Note that only 19% of bins that showed detectable change of H3K4me1 were included in the analysis. The other 81% of bins were indicated as ‘N.D.’ in the figure. The data of all the bins were included (WG) for comparison. Y-axis shows the change of FIRE score for the indicated bins. **(H)** The Boxplot shows that change of H3K27ac signal is correlated with change of FIRE scors. Similar to Figure 4D, the 10-kb bins were categorized to 4 groups: I, no H3K27ac peaks in either cell type; II, with WT specific H3K27ac peaks only; III, with DKO specific H3K27ac peaks only; IV, with shared H3K27ac peaks. y-axis shows Log2 transformed fold change of FIRE score between WT and DKO cells.

**Figure S5.**
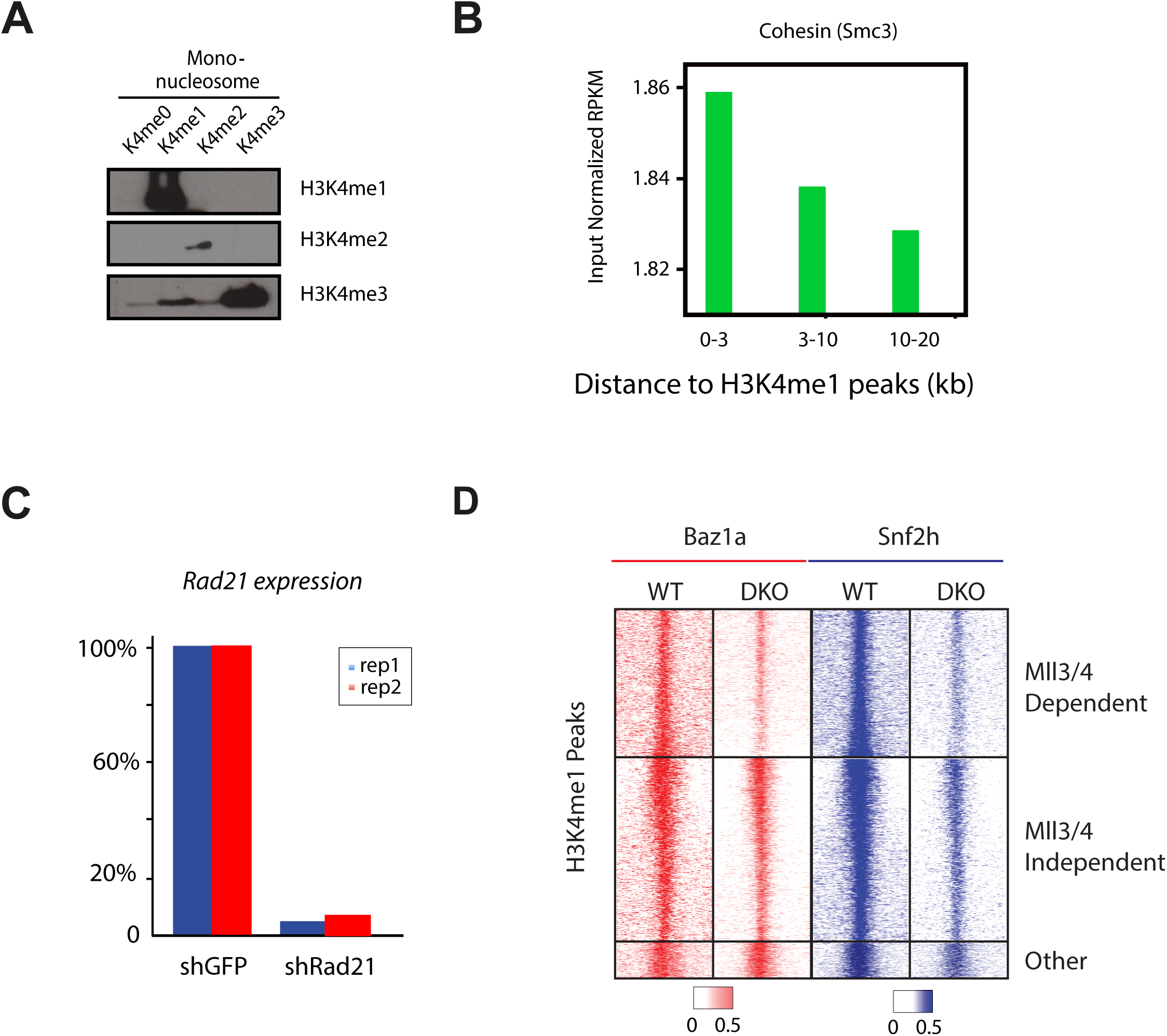
H3K4me1 Assisted Cohesin Binding to Chromatin, Related to Figure 3. **(A)** Gel images show the quality of mononucleosome assembly. Western blot shows that the assembled nucleosomes are modified specifically at lysine 4 of histone H3 (right). **(B)** Barplot shows that Cohesin/Smc3 is associated with Mll3/4 dependent H3K4me1 peaks. Cohesin is more enriched in the regions that are closer to Mll3/4 dependent H3K4me1 peaks than the distal regions. **(C)** qPCR confirms that Rad21 knock-down drops the mRNA level of Rad21 to less than 10% in WT cells. **(D)** Heatmap shows Baz1a and Snf2h binding around Mll3/4 dependent H3K4me1 peaks are dramatically decreased but only mildly affected around Mll3/4 independent H3K4me1 peaks or other H3K4me1 peaks. Each row show 10-kb bin centered by H3K4me1 peak summit.

**Figure S6.**
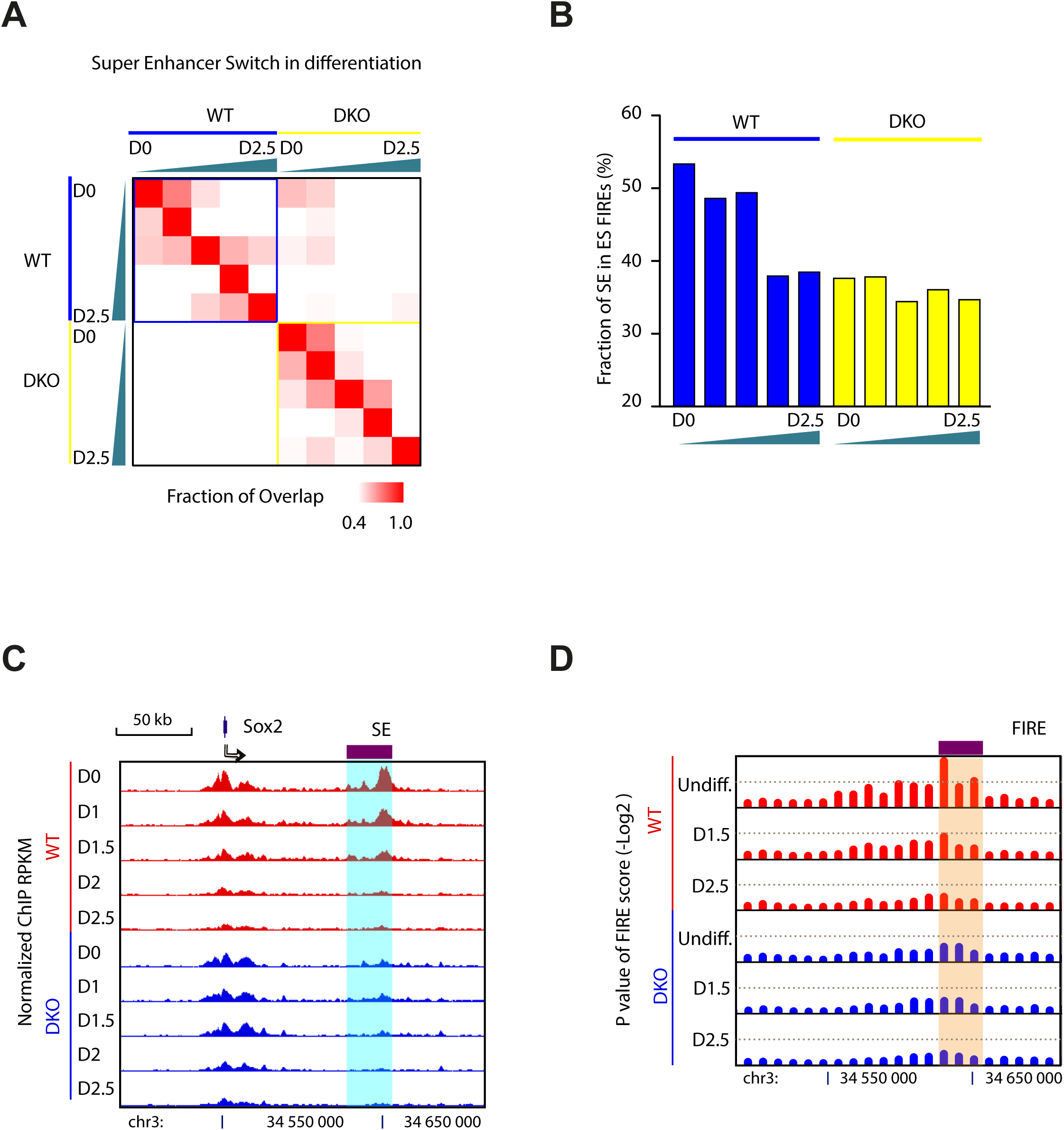
Super-enhancer Call Using H3K27ac ChIP-seq Data, Related to Figure 6. **(A)** Heatmap shows the overlapping of super-enhancers between cells at different time points during differentiation and between two cell types. Color key shows the fraction of overlapping. Green triangle shows the differentiation time, the thicker side representing later time points. D0, Day1; D2.5, Day2.5. Note that cells of the same type tend to show higher overlapping than between two cell types and that cells at consecutive time points show high fraction of shared super-enhancer. **(B)** Fraction of super-enhancer located in FIREs that are defined in WT cells. Note that over 50% of super-enhancers are located at FIREs and they are decreased along the differentiation. DKO cells show low overlapping of super-enhancers in WT FIREs confirming its different characteristics and cell identity. **(C)** Genome browser tracks show the change of super-enhancer at *Sox2* locus. Note that Sox2 super-enhancer is lost at Day 1.5. **(D)** Genome browser tracks show the change of FIRE at the same locus. Note that FIRE at Sox2 super-enhancer also gets lost at Day 1.5.

## ACKNOWLEDGMENTS

We would like to thank the Ren Lab members Drs. David Gorkin, Tingting Du, Miao Yu, Jason Guoqiang Li, as well as Drs. Shicai Fan (UCSD) and Xi Wang (DKFZ, Germany) for comments during manuscript preparation. We are also very grateful to Samantha Kuan and Dr. Bin Li for technical assistance. We thank UCSD Neuroscience Microscopy Shared Facility (NS047101) for FISH imaging and the Murre Lab (UCSD) for sharing protocols and discussions for FISH assay. This work was supported by the Ludwig Institute for Cancer Research (B.R.), NIH (P50 GM085764-04, 1U54DK107977-01) (B.R.), and an International Postdoc fellowship from the Swedish Vetenskapsrådet (537-2014-6796) (J.Y.).

## ACCESSION NUMBERS

Sequencing data have been deposited in Gene Expression Omnibus (GEO) under accession number GSE74055 (For the reviewing purpose, the following link should be used: http://www.ncbi.nlm.nih.gov/geo/query/acc.cgi?token=ctonkmiollmflir&acc=GSE74055).

## AUTHOR CONTRIBUTIONS

J.Y., S.A.C. and B.R. conceived the project. J.Y, S.A.C, A.L, T.L., A.Y.L., C.M.R., H.I., C.W., K.G., S.P. and Z.Y. carried out experiments. J.Y., I.J., R.F., M.H. and Y.Q. performed data analysis. J.Y., S.A.C. and B.R. wrote the manuscript. All authors edited the manuscript.

